# Spatio-temporal patterns of multi-trophic biodiversity and food-web characteristics uncovered across a river catchment using environmental DNA

**DOI:** 10.1101/2021.07.20.450136

**Authors:** Rosetta C. Blackman, Hsi-Cheng Ho, Jean-Claude Walser, Florian Altermatt

## Abstract

Accurate characterisation of ecological communities with respect to their biodiversity and food-web structure is essential for conservation. However, combined empirical study of biodiversity and multi-trophic food-webs at a large spatial and temporal resolution has been prohibited by the lack of appropriate access to such data from natural systems. Here, we assessed biodiversity and food-web characteristics across a 700 km^2^ riverine network over seasons using environmental DNA. We found contrasting biodiversity patterns between major taxonomic groups; local richness showed seasonally dependent and statistically significant increases and decreases towards downstream positions within the catchment for fish and bacteria respectively, while invertebrate richness remained unchanged with increased downstream position. The local food-webs, formed by these taxonomic groups, also showed a variation in their structure, such as link density and nestedness, to both space and time, yet these patterns did not necessarily mirror those of biodiversity and functional feeding characteristics. In order to conserve species diversity as well as their functional trophic integrity of communities, patterns of biodiversity and food-web characteristics must thus be jointly studied, as our results suggest that they are not directly scalable to each other even at the same spatial and temporal scales.

## Introduction

The study of biodiversity patterns^1–3^ and the characterisation of food-web structures^4,5^ are essential, yet often disconnected elements of the study of ecology. Understanding these patterns is not only of fundamental interest, but also needed to predict stability, functioning and resilience of natural ecosystems and to bend the curve of biodiversity loss in the context of anthropogenic pressures and contemporary global change^6^.

Studies on biodiversity predominantly focus on analyses of α-, β- and γ-diversity, and possible underlying fundamental drivers of their spatial or temporal patterns^7^. Freshwater rivers are highly spatially structured systems^8–10^, in which theoretical and empirical studies have identified characteristic patterns of biodiversity for specific groups. For example, fish a-diversity has been found to increase with increasing catchment area (i.e., distance downstream)^11^, whereas headwaters often show high endemic bacterial species richness^12^. Aquatic invertebrate biodiversity has been widely studied and exhibits complex pattern with studies demonstrating both disproportionately high biodiversity found in headwaters compared to downstream^13^ and also the reverse: a significant increase in biodiversity linked to a greater catchment size^14^, which is generally a combination of opposing natural and anthropogenic drivers. However, group-specific biodiversity patterns have been mostly studied in isolation from one another, although all species are present within the same system and trophically interact with each other. Indeed, recent theoretical work shows that contrasting patterns driven by species’ resource competition are possible^15^. Therefore, to ensure optimal strategies for conservation and understanding of biodiversity patterns across different organismal groups, an ensemble approach integrating major taxonomic and trophic levels is crucial to reveal how species are linked through trophic interactions^16^ and how food-web structures unfold.

Trophic interactions and food-webs by definition encompass multiple groups of organisms. Individual freshwater food-webs are well-resolved^17^, and often exhibiting distinct features, such as highly nested structures^18^ and prevalent omnivory^17,19^. Nevertheless, food-web studies often have a localised perspective due to methodological limitations of sampling food-web interactions and organismal occurrence in a standardized and comparable manner across different places and organismal groups^20–22^. Due to the same reason, these studies also tend to focus on simple spatial and environmental gradients^23^ or temporal change^24^ when spatio-temporal influences should be considered in conjunction^25,26^. This is particularly problematic in freshwater riverine ecosystems, that are characterised by a high spatio-temporal structure: they exhibit characteristic spatial structures, and have strong seasonal variations driven by changing abiotic conditions^27^ and pronounced life-cycle changes of key taxa inhabiting these systems. The variation in dynamics over the course of a year remains a significant gap in our understanding of freshwater food-webs^17,28^.

To effectively conserve riverine biodiversity, we must encompass spatio-temporal variation of multiple trophic levels to understand the underlying dynamics of both biodiversity patterns and food-web characteristics^4,25,26^. In particular, molecular monitoring techniques may now provide a suitable solution to break through the above-mentioned methodological constraints caused by sampling based primarily on species sight or capture. Environmental DNA, or eDNA, is the collection of DNA extracted from an environmental sample such as water, air or sediment^29^. By collecting eDNA we can screen samples for multiple taxonomic groups via metabarcoding^30^, thereby creating a biodiversity assessment suitable for food-web reconstruction.

Here, we use eDNA metabarcoding to assess patterns of biodiversity and reconstruct local food-webs via a metaweb-based approach^31^. We used high-throughput sequencing and a multi-marker approach to examine the occurrence of three taxonomically relevant groups, namely fish (via the 106 bp fragment of the 12S barcode region), invertebrates (a 313 bp fragment of the cytochrome *c* oxidase I region, COI) and bacteria (a 450 bp fragment of the 16S barcode region) in a large-scale river network over the course of three seasons (spring, summer and autumn). A metaweb details all possible interactions of those species found in a dataset, and then each local food-web is inferred from the subset of those species and interactions found at each site. The metaweb approach is suitable to capture food-web patterns based on taxa co-occurrence data in both terrestrial and aquatic systems^24,32–35^. This allowed us to test for association of biodiversity and food-web structures with network location, season, and the interaction between these two factors. We found contrasting effects on biodiversity patterns and food-web structure from spatial and temporal influences and their interaction, providing insight into the underlying changing ecosystem dynamics and indicating that effective and targeted conservation measures must consider them jointly.

## Results

### Data collection and community construction

We collected water samples from the upper Thur catchment, Switzerland, which covers approximately 700 km^2^ and is made up of three sub-catchments: River Thur, Necker and Glatt. Seventy-three field sites were selected to allow for maximum coverage across the catchment area, comprising of a broad size range of upstream drainage sizes (i.e., the drainage area size indicates the location of the site within the catchment: upstream sites having a small drainage area and lowland sites having a larger drainage, see Supplementary Fig. S1). Water samples were filtered on site, sealed with luer fittings, and placed in individual sealed bags in a cool box for transport to the lab, where they were stored at −20 °C until further processing. DNA extraction was performed in a specialist clean lab environment at Eawag, Switzerland and subsequent library preparation followed a uni-directional lab processing protocol where PCR product was not returned to the cleaned lab. A total of 12.29 million, 14.32 million and 14.02 million raw reads were produced from the 12S, COI and 16S libraries, respectively. After bioinformatic processing the average (±sd) sequencing read depth per sample was: 42,189 ±19,057, 46,573 ±407,659 and 24,646 ±27,612 for 12S, COI and 16S, respectively (see Methods and Supplementary Table S1 and bioinformatics workflow). For further analysis, ZOTUs were merged to genus level and only fish genera, invertebrate genera with an aquatic life stage and bacteria associated with freshwater were kept for further analysis (see Methods for further details). Analysis was performed on presence/absence data to merge the three libraries for the complete freshwater community and exclude a possible influence of uneven sample read depth generated from multiple markers (see also ^36^).

To quantify biodiversity, we calculated α-diversity (local richness) at genus level at each site and compared β-diversity (variation in community composition between sites) by using Jaccard dissimilarity. Jaccard dissimilarity was also partitioned into taxon replacement (turnover) and taxon loss (nestedness) components to assess the mechanisms contributing to the variation in community assemblage across the catchment^37^.

To measure structural food-web and functional characteristics of communities, we constructed a metaweb based on known interactions of genera classified into different functional feeding groups (Fig. 2; see also Methods and Supplementary Table S2 and S3). We defined a local food-web at each field site based on genera co-occurrence and the corresponding subset of interactions from the metaweb. With this approach, variation in food-web structural and functional characteristics emerge from the spatio-temporal differences in genus and thus feeding-group composition, which then determines broad changes in food-webs over time and space (See Methods for further details).

**Figure 1:**
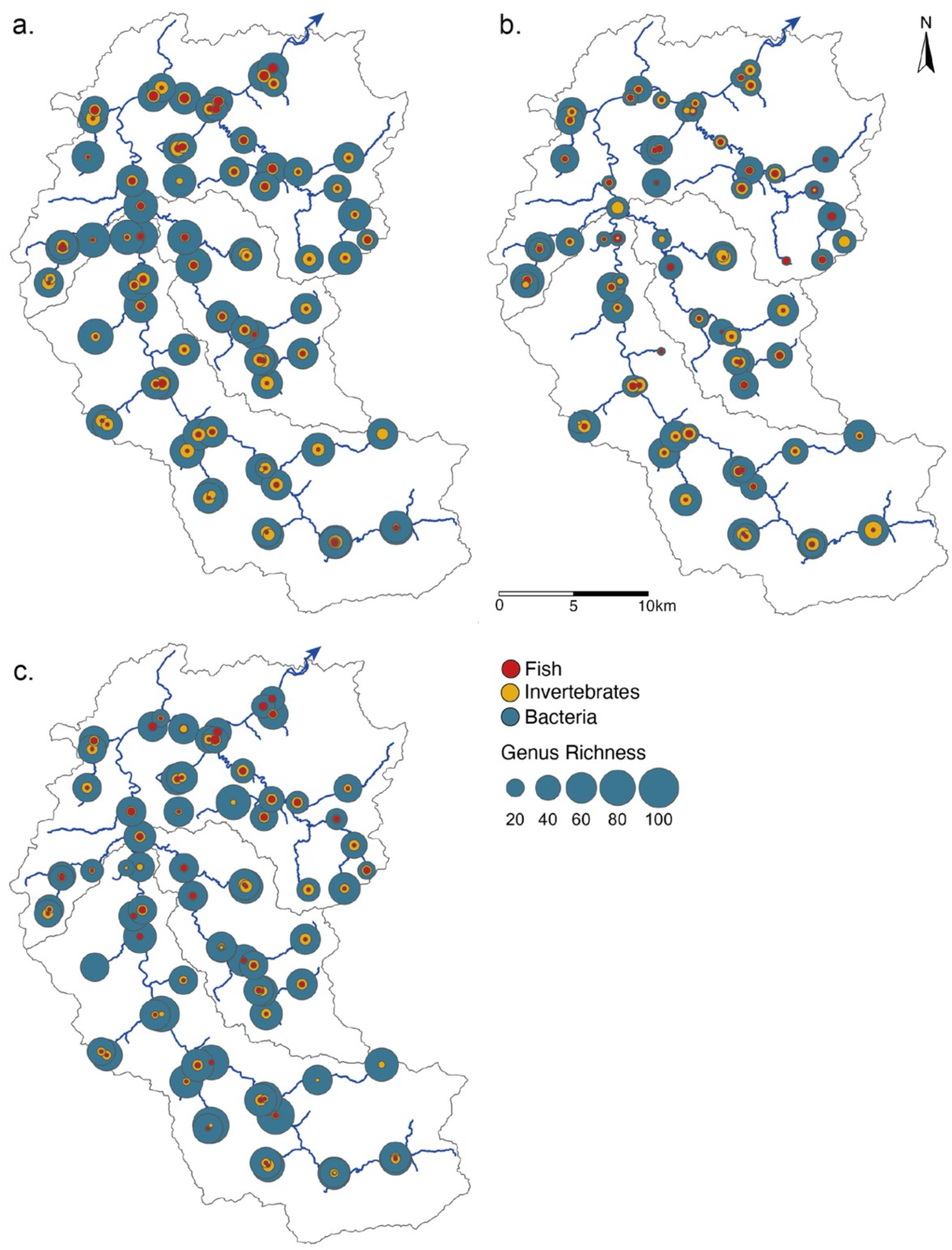
Genus α-diversity of bacteria, invertebrates and fish collected from eDNA samples in the Thur catchment in (a) spring, (b) summer and (c) autumn. The circles are scaled proportionate to the α-diversity of each group. Grey lines give the sub-catchments Glatt, Necker, and Thur, respectively and the blue arrow shows the direction of flow.

**Figure 2:**
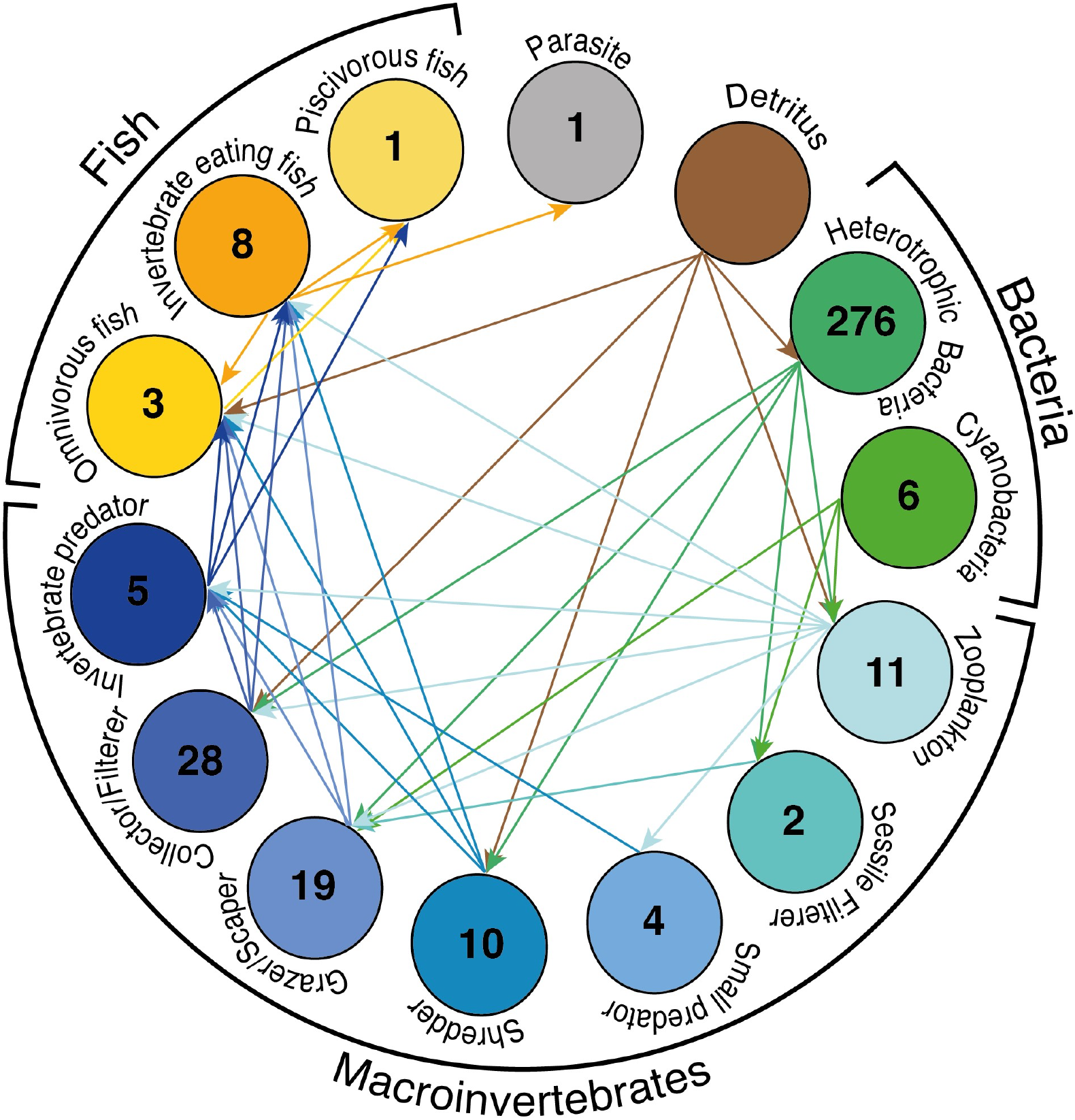
Trophic metaweb generated from eDNA data: All genera were categorised into feeding groups dependant on their dominant feeding behaviour, we then used this information to construct a trophic interaction metaweb using these links. Number of genera in each group is shown in circles, arrows indicate energy flow.

**Figure 3:**
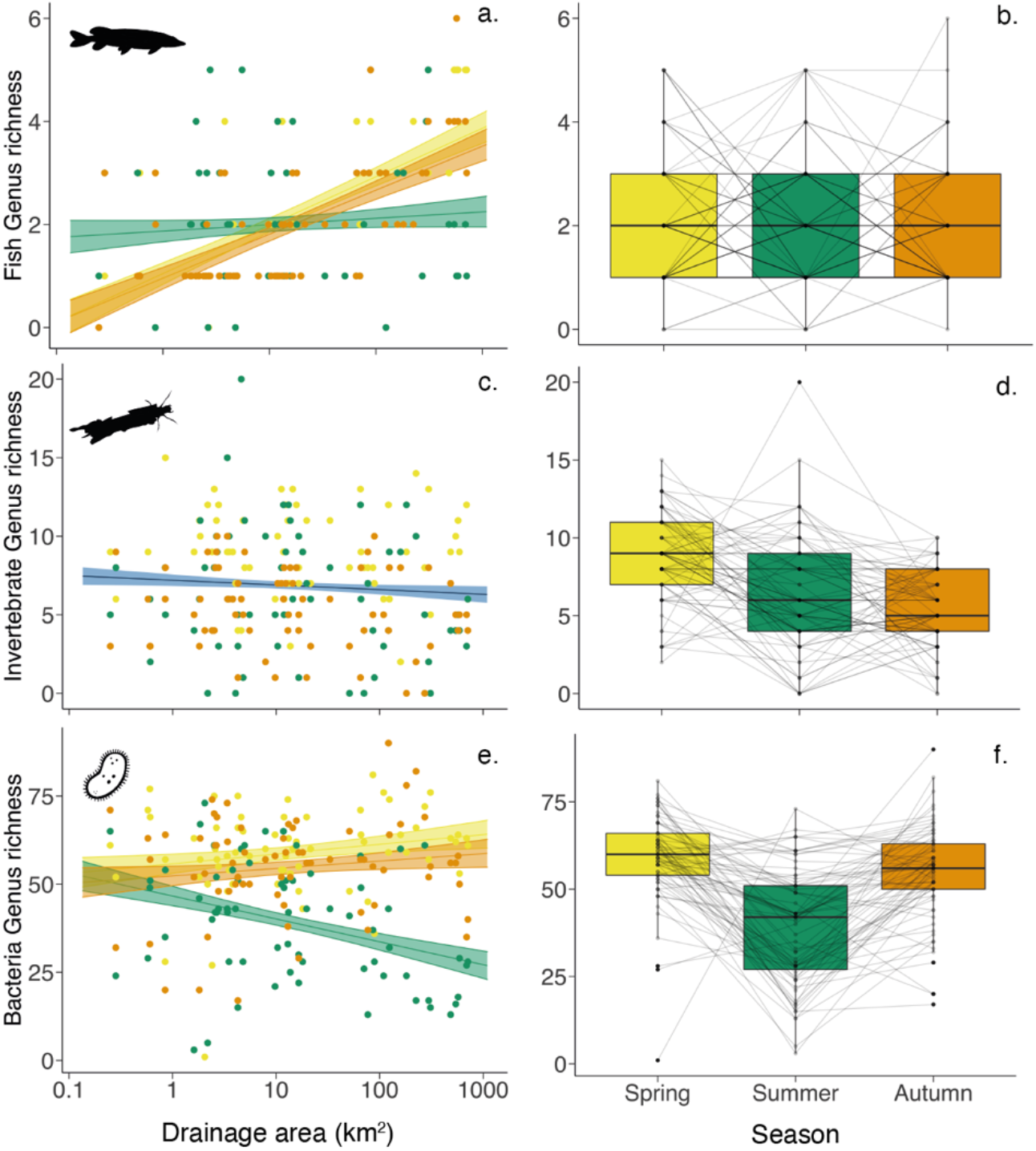
Genus α-diversity in space and time. Plots a, c and e show α-diversity in each group as a function of drainage area. Lines indicate linear mixed-effects models with shaded area showing 95% confidence intervals as calculated using the model predictions and standard error. Plots a and e show the models with a significant interaction between drainage area and season with the difference across each season (colour represents season). Plot c shows no significant effect of this interaction, the line indicates the model output of all data as a function of drainage area which is not significant (blue). Plots b, d and f show change in α-diversity in each group over season with samples sites linked by grey lines over the three sampling seasons. Colour represents season: yellow – Spring, green – Summer and orange – Autumn.

To assess the relationship between α-diversity of each group (fish, invertebrates and bacteria), food-web structural and functional characteristics with site location within the catchment, we conducted linear mixed-effects model analyses. Drainage area (km^2^) was log transformed to aid model assumptions for normality. For each dependant variable, drainage area and season were the fixed effects, while site was the random effect. We ran models with and without an interaction of the two fixed effects to determine the overall effect of factors, and report below the results based on the interaction models (see supplementary Table S5). As post hoc contrast testing, we ran estimated means of linear trends to compare the slopes from the mixed-effects models and estimated marginal means of data over season only to determine the influence of seasonal changes on all α-diversity and food-web measurements (see Methods for further details). To examine the effect of river distance on β-diversity we constructed a matrix of pairwise distances for sites that were connected along the fluvial network and to examine the effect of river distance on β-diversity we performed a Mantel test (see Methods and Supplementary Table S11).

### Spatial and temporal biodiversity patterns

In total we detected 374 genera across three organismal groups associated with freshwater, including 12 fish genera, 80 invertebrate genera and 282 bacteria genera. When combining all seasons, α-diversity (genus richness) ranged between 8–96 genera with all taxonomic groups combined (Fig. 1). Over the different seasons, mean local α-diversity was 70 (range 10–92) in spring, 48 (range 8–85) in summer, and 63 (range 19–96) in autumn (Supplementary Fig. S2 and Table S4). We used mixed-effects models to assess the influence of drainage area and season on the local α-diversity of each group (Fig. 3, for detailed model output see Supplementary Table S5–S8). Of the three taxonomic groups, interaction between drainage area and season was found to be significant for both fish and bacteria α-diversity (p < 0.001 for fish and bacteria, Table S8), in this study this indicates that the effect of drainage area is modulated by the season rather than vice-versa. By examining the slopes generated from the mixed-effect models for each season, the effect of drainage area differs across the seasons in both fish and bacteria α-diversity (Fig. 3a and e, Supplementary Table S9). Fish α-diversity in summer has a flatter slope and is significantly different compared to spring and autumn (p < 0.001 for both) both with steep increases in richness with increase of drainage area (Fig. 3a). Bacteria α-diversity also showed a similar trend such that summer diversity spatial pattern is significantly different from both spring and autumn sampling seasons (p < 0.001 and p < 0.01, respectively; Supplementary Table S9); however, this is mostly because of a sharp decline in richness with increased drainage area in summer (Fig. 3e). Examining the mean α-diversity of genus across seasons (i.e., collapsing the spatial axis) shows no significant change in fish (mean number of 2 genera and range 0–6, Fig. 3b), but for bacteria we find a significantly lower α-diversity in summer compared to spring and autumn (p < 0.001 for both, Fig. 3f and Supplementary Table S10). Looking at invertebrate α-diversity we do not see a significant interaction of drainage area and season (p = 0.295, Fig. 3c and Supplementary Table S5), but season as a significant main effect on it (p < 0.05, Supplementary Table S8). Examining the means across the seasons further shows a significant drop in α-diversity of invertebrates from spring to summer (p < 0.001) and summer and autumn (p< 0.0001, Fig 3d and Supplementary Table S10).

Regarding the analysis of β-diversity across the catchment, the Jaccard’s dissimilarity significantly increased with increasing pairwise river distance among sites in all organismal groups across all seasons, apart from bacteria in spring and autumn (fish Mantel statistics Spring: 0.143, p < 0.01, Summer: 0.171, p = 0.001, Autumn: 0.321, p = 0.001; invertebrate Mantel statistics Spring: 0.095, p < 0.05, Summer: 0.169, p = 0.001 and Autumn: 0.114, p < 0.05; bacteria Mantel statistics Spring: −0.058, p = 0.834, Summer: 0.179, p < 0.01 and Autumn: 0.069, p = 0.12, See Supplementary Table S11). Further partitioned analyses on taxon replacement and loss revealed contrasting patterns. Taxon replacement between sites increased over river distance for all groups in most seasons (significant or marginally significant, see Supplementary Table S11), apart from fish (Mantel statistics: 0.07, p = 0.891) and bacteria (Mantel statistics: 0.016, p = 0.34) in spring. In contrast, taxon loss was only found to significantly increase over river distance for fish in spring and autumn (Mantel statistics: 0.219, p = 0.001 and 0.059, p = 0.05, respectively) and bacteria in summer (Mantel statistics: 0.089, p < 0.05).

### Spatial and temporal changes in food-web structural and functional characteristics

We examined commonly used food-web structural characteristics (link density, connectance, nestedness, level of omnivory, coherence, number of links, modularity, and robustness), as well as functional characteristics (functional diversity and redundancy, Fig. 2 & Supplementary Fig. S2 and Table S3). For food-web structure, here we describe the results of link density, connectance, nestedness and level of omnivory (hereinafter, omnivory) as the most ecologically important food-web descriptors in our study, while the results from the remaining descriptors are presented in the Supplementary Information (Table S5–10 and Fig. S4). As with the biodiversity patterns, we find a significant interaction between drainage area and season for connectance, nestedness, and marginally for omnivory, but not for link density (p < 0.05, p < 0.001, p = 0.066 and p = 0.138, respectively; Fig 4 and Supplementary Table S8). Food-web connectance, nestedness and omnivory increased with increasing drainage area, particularly in summer, and in contrast showed a decreasing trend in spring and autumn (Fig. 4c, e, and g). There, summer patterns were significantly or marginally different from the ones of autumn (p < 0.05, p < 0.001, and p = 0.074, respectively; Supplementary Table S9). Whereas link density showed only a marginally significant decrease with drainage area (p = 0.064; Supplementary Table S8) and a significant effect of season (p < 0.05; Fig. 4a and Supplementary Table S8). When we examined mean food-web measurements across the seasons, link density was significantly higher in spring than in summer and autumn (p < 0.001 for both, Fig. 4b and Supplementary Table S10) whereas connectance and nestedness were significantly lower in autumn than in spring and summer (all p < 0.01 or 0.001, Fig. 4d and f and Supplementary Table S10), and omnivory was significantly higher in summer than in autumn (p < 0.001, Fig 4h and Supplementary Table S10).

**Figure 4:**
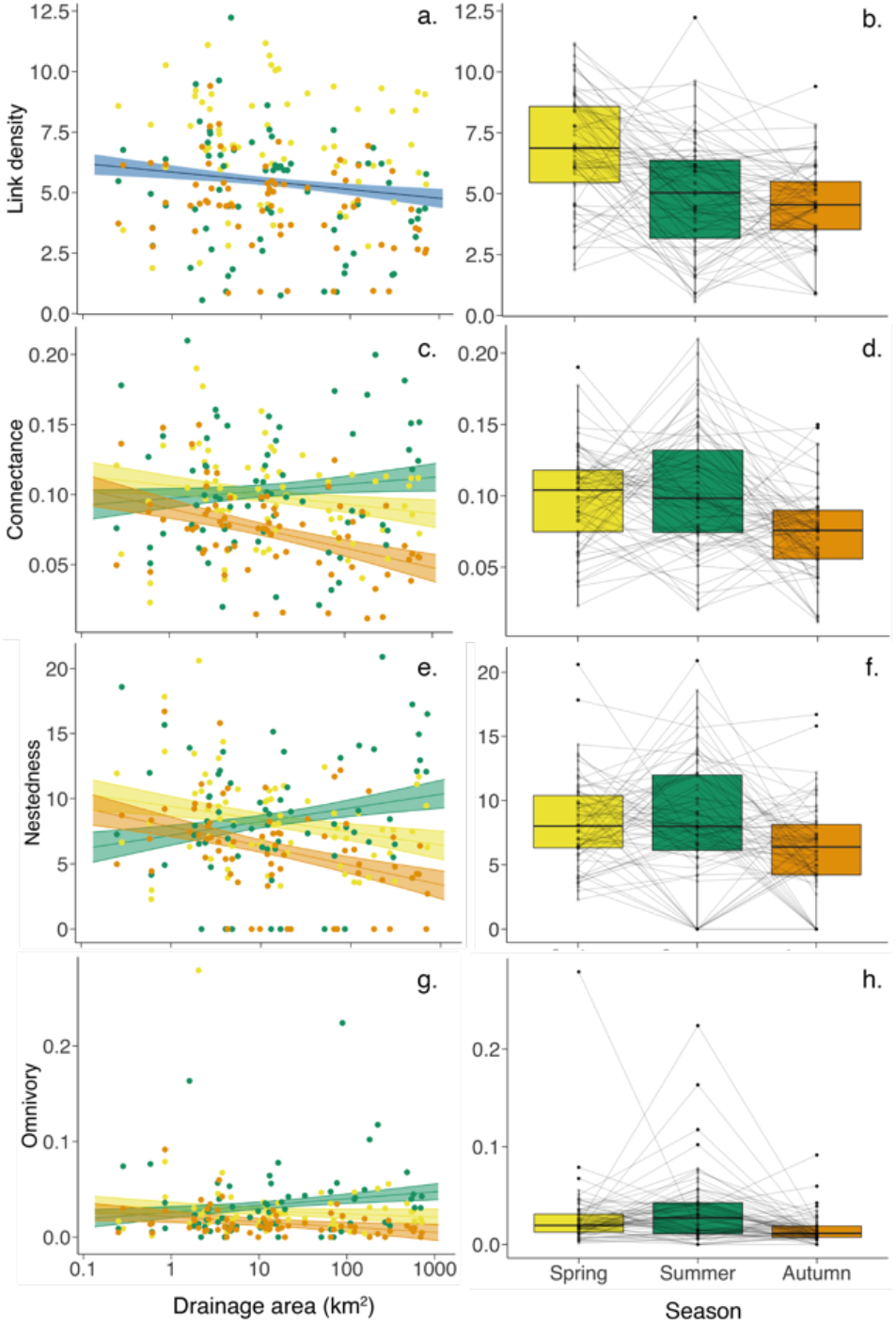
Food-web structural characteristics: Plots show food-web structural characteristic: a and b: Link Density, c and d: Connectance, e and f: Nestedness and g and h: Omnivory. Lines indicate linear mixed-effects models with shaded area showing 95% confidence intervals as calculated using the model predictions and standard error. Plots a shows no significant interaction between drainage area and season, therefore the line indicates the model output of all the data as a function of drainage area which is marginally significant (blue). Plots c, e and g show the models with a significant interaction with the difference across each season (colour represents season). Plots b, d, f and h show the change in characteristic over season with samples sites linked by grey lines over the three sampling seasons. Colour represents season: yellow – Spring, green – Summer and orange – Autumn.

Functional diversity showed no significant interaction between drainage area and season (p = 0.354, Supplementary Table S8), but a marginally significant decrease with increasing drainage area while exhibiting no change with season (p < 0.05 and p = 0.147, respectively; Fig. 5a and b and Supplementary Table S8). Functional redundancy showed a significant interaction between drainage area and season (p < 0.01, Supplementary Table S8). Across seasons, the mean functional redundancy was significantly lower in summer than in autumn (p < 0.05, Fig. 5d and Supplementary Table S10).

**Figure 5:**
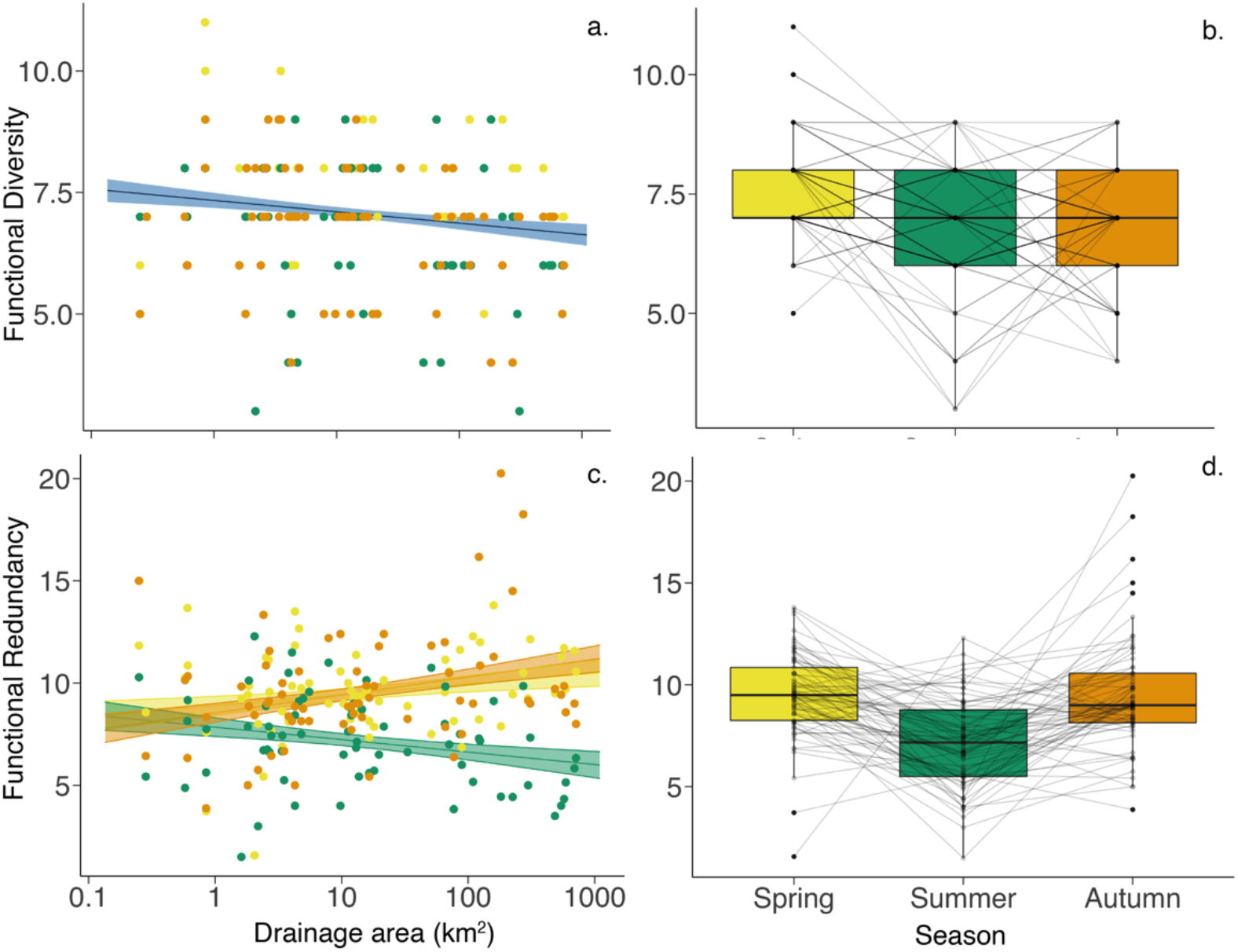
Functional diversity and functional redundancy. Plots show functional dynamics: a and b: functional diversity and c and d: functional redundancy. Lines indicate linear mixed effect models with shaded area showing 95% confidence intervals as calculated using the model predictions and standard error. Plot a shows no significant effect of the interaction between drainage area and season therefore the line indicates the model output of all the data as a function of drainage area which is marginally significant (blue). Plot c shows the model with a significant interaction with the difference across each season (colour represents season). Plots b and d show the change over season with samples sites linked by grey lines over the three sampling seasons. Colour represents season: yellow – Spring, green – Summer and orange – Autumn and blue – all seasons combined.

## Discussion

Studies of biodiversity and food-web assemblages in riverine networks are often constrained to local scales or aggregated to a single time point, which in essence fails to capture the spatial processes and the temporal fluctuations that together play a key role in community dynamics present within a river network^8^. Our study is one of the first assessments utilising data derived from eDNA metabarcoding to describe biodiversity patterns and food-web characteristics^38–40^ in a whole river network at a spatially and temporally large scale. By taking this approach, we were able to detect contrasting group-specific spatial and temporal biodiversity patterns, as well as food-web structural and functional characteristics whose spatial responses also showed strong dependence on the season, yet in different ways. Overall, our findings support the need to include multifaceted biomonitoring, i.e., at taxa and community levels and regarding richness as well as biological interactions, conducting across both spatial and temporal scales, to better understand changes in ecosystems^41^. Such a comprehension is particularly crucial as we are facing contemporary global change and biodiversity loss.

Aquatic biodiversity is subject to fluvial influences within a dendritic network, whereby spatial patterns of α-diversity and β-diversity are known for key groups, such as fish, invertebrates and microbes^11–14,42^. Our data is congruent with previous studies on fish diversity^11,36^, in that both α-diversity significantly increased downstream, and β-diversity (community dissimilarity) significantly increased with river distance in all three seasons. Bacterial richness was highest at the top of catchment, which was also in line with previous studies^37^, albeit only in the summer sampling campaign. Similarly, bacterial β-diversity also increased with river distance during this season. We therefore postulate that bacteria are able to disperse through the focal catchment during higher flows (spring and autumn), which makes bacterial spatial diversity patterns subject to seasonal influences. Unlike the former two groups, invertebrate richness exhibited no significant influence of drainage size but a seasonal change. The seasonal variation in invertebrate diversity can be linked directly to the emergence of the non-aquatic adult life stage of several macroinvertebrate genera detected (Diptera, Ephemeroptera, Plecoptera and Trichoptera), which takes place in the late spring and summer months^27^, and literally removes these organisms from the aquatic communities as they become air bound. For the lack of spatial effect, this result is contrary to previous studies which have shown clear, although variable, spatial patterns for invertebrate communities in river systems^13,14,43^. As previous studies often looked at spatial scales much larger or much smaller than our, this disparity may be due to a mismatch in scale looked at here, or reflect different strengths of anthropogenic drivers that may have opposing effects. For example, Finn and colleagues^13^ suggested headwaters harbour disproportionately high invertebrates compared to lower points in the catchment, but their studied focused on small (1–2) and mid (3–4) stream orders only, whereas the sites in our study ranged from small to much larger rivers (stream orders 1–7, ^44^). Contrastingly, when looking at scales about 50-fold larger, Altermatt and colleagues^14^ showed the number of key aquatic invertebrate taxa (Ephemeroptera, Plecoptera and Trichoptera) increased with catchment size, however they also found a combination of local factors (catchment areas, drainage area, elevation and network centrality) had the greatest influence on local invertebrate α-diversity. Possibly, the scale at which we study invertebrate α-diversity and contributing local factors falls between such small-scale vs. large-scale perspective, such that we detected regional patterns where α-diversity remains relatively constant throughout the catchment. These findings thus expand on past studies, highlighting the influences of spatio-temporal interactions on freshwater biodiversity, and the importance of choosing a scale at which we monitor biodiversity patterns.

By resolving the fundamental trophic relationships among broad feeding groups, we established a trophic interaction metaweb for the three taxonomic groups examined in this study. The metaweb with genus co-occurrence data thus allowed our investigation of both spatial and temporal influences on the characteristics of local food-webs. With regard to food-web structure, we found a significant interaction between drainage area and season that associated with connectance, nestedness and omnivory, whereas link density was influenced in isolation significantly by season and marginally by drainage area. Overall, the impact of season turns out to be prevalent in all four focal structural characteristics, highlighting the importance of considering seasonality of food-web structure. By definition, a lower link density indicates a higher proportion of trophic specialist (who interact with relatively fewer other taxa) in the food-web. Therefore, combining observed link density and α-diversity patterns together, reveals that communities are larger and have more specialists toward downstream in wetter seasons (spring and autumn), while smaller and fewer generalists are found towards downstream in the dry season (summer). Such changes are likely driven mainly by bacteria given their dominance in the communities. Thus, the unique summer spatial patterns (in comparison to spring and autumn) for food-web connectance, nestedness and omnivory may reflect the similarly unique bacterial α-diversity summer spatial pattern. Temporally, all four focal food-web structural characteristics were lowest in autumn, despite overall α-diversity being lowest in summer. This indicates that the seasonal variation in food-web structure found in this study does not merely reflect the overall richness (i.e., community size) change, but rather the change in genera and thus feeding-group composition. Previous studies on freshwater food-webs that have examined temporal changes have often found reduced productivity as the main driver behind a declined food-web structure in winter vs. summer^20,45^. Thus, the restrained food-web structural characteristics we see in autumn may capture the start of the productivity restriction seen in food-webs over the winter months^46^, allowing only some the feeding groups to persist (as further suggested by our functional redundancy results, see below). Indeed, the detected less-connected and less-nested food-web structure also implies weaker resource competition among consumers, which may be necessary for their coexistence in the food-web^47^, especially when competition becomes more costly with lower resource productivity in autumn.

Moving from broad food-web patterns to examining feeding groups enables us to study fundamental functional changes within the community and possible effects at the ecosystem level^17^. We find a marginally significant effect of drainage area on the functional diversity, which is highest upstream. Such a spatial pattern along the hierarchical river network is congruent with to the River Continuum Concept (RCC), which states increased diversity of invertebrate functional feeding groups at low stream orders^48,49^ (i.e., upstream). In river systems allochthonous material (i.e., the dominant organic material) in forested headwaters is processed by specialist shredder, collector/filterers, and grazer/scrapers, while downstream autochthonous organic material is utilized by fewer functional feeding groups^48^. Therefore, the pattern we observed is to be expected and shows the suitability of our approach of using single overarching functional feeding groups (i.e., a single dominant functional feeding group per genera), which simplified the life cycle and changes in feeding behaviour observed in invertebrates (the dominant group within the functional feeding groups). Interestingly, the functional redundancy we observe gives us more information about the changes exhibited at a community level. Functional redundancy, i.e., genera per functional feeding group, was interactively influenced by drainage area and season, showing a unique decrease toward downstream in summer, contrasting the opposite spatial trend in spring and autumn. Also, the functional redundancy was lower in summer than in autumn. Since this measurement accounts for the number of genera, its spatio-temporal patterns could again largely reflect the variation in bacteria richness. Nonetheless, the particularly high functional redundancy in autumn suggests that, although fish and bacteria richness increase toward downstream, some key invertebrate taxa of well-connected feeding groups (i.e., interact with many other groups) can become absent particularly in this season, and losing such feeding groups cause a constriction of local food-web structure as we observed. Furthermore, although both the food-web structural and functional characteristics are determined by genera occurrence, their inconsistent and variable temporal patterns imply that multiple feeding groups share similar trophic roles in the food-web (Fig. S3), though they each adopt specialised feeding behaviour within the catchment and thus perform distinctive ecological functions (e.g., shredders, collectors, filterers). In other words, the structure and function of a food-web does not necessarily match and synchronise (*sensu stricto*^22,50^). These results are particularly encouraging, because the patterns of both food-web structure and functional diversity are known to be important for ecosystem health assessment and identifying potential vulnerability to perturbation^17,26,51^. Addressing the consistency of their patterns across spatial and temporal scales will likely lead to novel and comprehensive understanding of biodiversity and ecosystem function loss due to environmental change.

In our study, the functional feeding groups were defined at the level of genus, while we expect investigations at finer resolutions will be promising for future work that can reveal not only more accurate patterns, but also the influences of sampling taxonomic resolution. Similarly, our selection of focal taxa may have influenced the food-web patterns we detected. With the selected three key taxonomic groups in riverine ecosystems, we present a broad and relevant view capturing trophic roles from basal resources (cyanobacteria) to top consumers (piscivorous fish), captured by three relatively broad metabarcoding markers. However, we also by default excluded some further groups, such as algae or terrestrial taxa, which could have been relevant as primary producers and as terrestrial-aquatic linkages, respectively. For example, it is known that algae become an increasingly important resource when moving from allochthonous inputs in headwaters to larger streams with increased light levels further down the catchment^48,49^. The choice of taxonomic groups looked at was both methodologically defined, as well as driven by the goal to have an overseeable and clearly aquatic-focused view on food-webs. Thereby, we here capture the spatio-temporal patterns of an (dominant) subset from the real-world communities and food-webs in which even more species are involved.

By using the eDNA technique, we gain three notable advantages: its scalability for monitoring complex and large systems, its reusable nature, and its being a non-invasive method of collecting biodiversity information^41,52^. Our understanding of how the information we ascertain from eDNA sample collection has greatly improved in recent years due to studies on the hydrological influences^53^ and a general understanding of the rate of eDNA persistence in lotic ecosystems^54^. However, the successful detection of taxa with eDNA is also linked to the ecology of individual species^54^, and some seasonal variation in the detection of several taxonomic groups is known^55–58^. Therefore, it is possible that some taxa that were not detected in the colder season (autumn) in this study were false-negative records. However, these non-detections are likely linked to low abundance or low metabolic rates, and thus these species are, while not physically absent, at least “relatively absent” in ecological terms.

In summary, we reconstruct comprehensible multi-trophic community and evaluate their characteristics at a large spatial scale (i.e., the catchment) and over time (i.e., seasons), by exploiting broad species occurrence information derived by eDNA sampling and combining multiple markers. For both biodiversity and food-web characteristic, we detect spatio-temporal patterns, and those of the former are different across taxonomic groups while the latter across measurements. Specifically, we identify inconsistent patterns between food-web structural and functional feeding characteristics, as well as a reduction in local food-web structure particularly in autumn. Our approach provides a first demonstration of the application of eDNA to a complex river network for the monitoring of not only biodiversity but also food-web patterns, paving a way for establishing long-term, repeated monitoring of complex communities, and potentially also in other systems worldwide. Biodiversity in freshwater systems face huge threats from anthropogenic pressures, including global climate change. To protect and conserve the systems, it is essential to establish vital information on the changes in biodiversity and food-web composition over spatio-temporal scales for the detection of these stressors.

## Methods

### Site selection and eDNA sample collection

Environmental DNA water samples were collected from the edge of the waterbody and filtered on site using single use disposable 50 ml syringes. At each site the syringe was filled and refilled 10 times from the river and the water was filtered through a 0.22 μm sterivex filters (Merck Millipore, Merck KgaA, Darmstadt, Germany). Two syringes and two sterivex filters were used per site, equating to 1 L of samples water per site. The filters were then sealed with luer caps, placed in individually labelled bags in a cool box for transport to the lab, where they were stored at −20 °C until further processing. Sampling was carried out over five consecutive days in May (spring), August (summer) and October (autumn) in 2018. During the summer sampling campaign, four sites were dry and could not be sampled (total samples across all seasons is n = 215). Negative field control samples were collected on each sampling day by filtering 500 ml of ddH2O through a sterivex filter in the same way as field samples were collected (n = 15). Negative field samples were processed alongside field samples. All samples were placed in labelled bags and cooled in a cool box until frozen at −20 °C on return to the laboratory.

### Extraction and library preparation

DNA extraction and first round PCR set up was carried out in a specialist eDNA clean lab, with separate lab facilities for post PCR workflows. Samples were extracted using DNeasy PowerWater Sterivex Kit (Qiagen, Hilden, Germany) following the manufacturer’s protocol. Prior to extraction, each sterivex was defrosted at room temperature and then wiped with 10% bleach, then 70% ethanol solution prior to extraction to remove any DNA from the outside of the filters. Extraction controls were carried out alongside sample extractions and consisted of blank sterivex filters (n = 5). Samples selected at random were analysed using a QuBit 3.0 fluorometer for double stranded DNA concentration, values measured between 0.317 - 27.5 ng/μl. All negative controls (field and extraction) were tested and recorded below detection limits.

Sample replicated were pooled for subsequent sequencing with the following markers: a 106 bp fragment of the mitochondrial 12S marker (^59^, hereafter referred to as 12S) used to amplify vertebrate DNA, a 313 bp fragment of the mitochondrial cytochrome oxidase I marker (^60^ and ^61^, hereafter referred to as COI) used to amplify metazoan DNA and a 450 bp fragment of the V3-V4 region of the 16S marker (^62^, hereafter, 16S) used to amplify bacteria and archaea DNA (See Supplementary Information Table S8 for primer sequences). Positive controls (n = 6 per library prep, see Supplementary Information Table S9) and PCR negative controls (2 μl of ddH2O, n = 11 per library prep) were included in each library. Each library consisted of 252 samples in total (including positive and negative controls).

Library preparation followed a two-step PCR process for both the 12S and COI markers, the 16S library was carried out using a three-step PCR^63^ (See Supplementary Information Methods for full details), all samples were amplified in triplicate. After the initial amplification, where Nextera® transposase sequences (Microsynth, AG, Balgach, Switzerland) are added to the PCR product, all samples were tested for amplification success with the AM320 method on the QiAxcel Screening Cartridge (Qiagen, Germany). PCR products were cleaned using ZR-96 Plate clean-up Kit (Zymo Research) following the manufacturers protocol with the minor modification by which the elution step was prolonged to 2 minutes at 4000 g. The clean amplicons were indexed using unique combinations of the Illumina Nextera XT Index Kit A, C and D following the manufacturer’s protocol (Illumina, Inc., San Diego, CA, USA). A reaction contained 25 μL 2x KAPA HIFI HotStart ReadyMix (Kapa Biosystems, Inc., USA), 5 μL of each of the Nextera XT Index adaptors and 15 μL of the DNA templates. The final PCR products were then cleaned using the Thermo MG Magjet bead clean up kit (Thermo Fisher Scientific Inc., MA, USA) and a customized programme for the KingFisher Flex Purification System (Thermo Fisher Scientific Inc., MA, USA) in order to remove excessive Nextera XT adaptors. The cleaned product of 50 μl was then eluted into a new plate and stored at 4 °C prior to quantification. For each library, the clean indexed amplicons were quantified in duplicate on a Spark Multimode Microplate Reader (Tecan, US Inc. USA) prior to equimolar pooling using the BRAND Liquid Handling Station (BRAND GMBH + CO KG, Wertheim, GE). Negative controls were used here to determine if any contamination had occurred at either the PCR1 or PCR2 stage. No such amplification was detected when the samples were run on the QIAxcel and the maximum amount of each negative sample was pooled with the other samples for library preparation. Library concentration was quantified on the Agilent 4200 TapeStation (Agilent Technologies, Inc., USA) and verified by the Qubit the HS Assay Kit. Paired-end sequencing was performed on an Illumina MiSeq (Illumina, Inc. San Diego, CA, USA) at the Genetic Diversity Centre at ETH (See supplementary information Table S1 for library loading information).

### Bioinformatics

After each of the libraries were sequenced, the data was demultiplexed and reads were quality checked using usearch v11.0.667^64^ and FastQC^65^. Raw reads were first end-trimmed, merged and full-length primer sites were removed using usearch v11.0.667^64^ (16S reads were merged using Flash v1.2.11,^66^). The merged and primer trimmed reads were quality filtered using prinseq-lite (0.20.4). The UNOISE3 (usearch v11.0.667) workflow with an additional clustering of 99% (usearch v11.0.667) identity was applied to obtain error corrected and chimera-filtered sequence variants ZOTUs. The ZOTUs were then clustered using a 97 %, similarity approach and taxonomic assignment with a 0.85 confidence threshold was performed using SINTAX in the usearch software v11.0.667^64^ with following databases: 12S: NCBI BLAST (v200416), COI: Custom reference database (Including MIDORI un-trimmed (V20180221) and 16S: SILVA (V128). See Supplementary Information Methods for all bioinformatic parameters and reference database information used for each library. Prior to downstream analysis, positive samples were examined from each library to see signs of potential contamination. Once we had removed terrestrial genera (such as *Bos taurus,* which was identified due to the use of BSA in the 12S library preparation), we applied a minor contamination of 0.1% to reduce errors caused by tag-jumping or minor contamination in library preparation^67^.

### Biodiversity patterns

We calculated α-diversity (genus richness) at each site and compared β-diversity between sites by using Jaccard dissimilarity using the betapart R-package^37^. We constructed a matrix of pairwise distances between sites along the fluvial network with the igraph R-package^68^ and compared the similarity between β-diversity (including further partition β-diversity into species loss and replacement) and river distance using the Mantel test with the vegan R-package^69^. Sites which did not record genus from a target group were removed from this analysis: 12S dataset: 4 sites were removed from Spring, Summer and Autumn, the COI dataset 4 sites were removed from the Summer and 3 from the Autumn analysis, no further sites were removed from the 16S Bacteria dataset.

### Feeding group assignment

Functional feeding groups were determined based on literature, species inventories and targeted expert knowledge^70,71^ for fish and invertebrates. Genera of bacteria were included if the phyla they belong to has a strong association with freshwater habitat^72^, bacteria were then broadly divided into heterotrophic and cyanobacteria, the latter constituting a basal resource. An assumptive constant basal resource of detritus was also included in all food-webs. Genera were designated into one of eleven feeding groups: Parasite, Piscivorous fish, Invertebrate eating fish, Omnivorous fish, Invertebrate predator, Collector/Filterer, Grazer/Scraper, Shredder, Small predator, Sessile filterer, Zooplankton, Cyanobacteria and Heterotrophic bacteria (see Supplementary Table S2 for descriptions of the feeding-group categories and Table S3 for assignment of genera to each feeding group).

### Food-web structure

We constructed a metaweb based on known trophic relationships among feeding groups (Supplementary Table S2), then defined local food-webs using this metaweb at each field site, based on co-occurrence of genera (nodes) and their interactions (links) (see Supplementary Fig. S5 for site food-web examples and associated GitHub repository for information on for feeding-group occurrence found in each sample). The metaweb approach has been applied to identify food-web characteristic change across spatial gradients in terrestrial ecosystems^32^, and temporal changes in individual aquatic ecosystem^24^, but has yet to be used at a large spatial scale in an aquatic ecosystem over time.

We then quantified common food-web structural characteristics for each local food-web generated. These included fundamental node-link composition features, i.e., number of links (L), link density (L/S, the number of feedings links per taxa divided by the genus α-diversity at each site) and connectance (L/S^2^, the proportion of realised interactions). To further explore the holistic topology of these food-webs and their change over space and time, we adopted the following indices to determine some more features. Nestedness is an indication that the diets of specialist taxa are subsets of generalist taxa’s diets, and was calculated using the nestedness metric based on Overlap and Decreasing Fill (NODF) function in vegan^69^. Modularity is indication of a less nesteded network^73^ in that the nodes are characterised into modules which, unlike nested structures, do not overlap, and was calculated using the multilevel.community() function from the igraph package^68^. Omnivory is defined as having a mixed trophic-level diet. Thus, the more omnivores a community has, the less coherent (see below) the food-web^74^. The level of omnivory is often debated as a measure of food-web stability, and weak omnivorous interactions are likely to lead to a more stable food-web^75^. Trophic level and level of omnivory of genera were calculated using the methods stated in William and Martinez^19^, and omnivory was averaged over all consumers in the food-web as a community-level index. Coherence is defined as the overall degree of homogeneity of the difference between trophic levels of every consumer resource pair within the local food-web. As described by ^74^, the coherence of a network can reliably be used to establish the stability of a network by looking at the distribution of trophic distances over all links in each network. For example, a perfectly coherent network (q = 0) implies that each taxon within the food-web only occupies a single trophic level. Coherence was calculated using the methods stated in ^74^. Robustness was defined by Dunne and Williams^5^ as the *“proportion of species subjected to primary removals that resulted in a total loss … of some specified proportion of the species”.* In our study, we used the commonly adopted 50% (i.e., R_50_) as the threshold proportion of species lost at which we evaluated robustness, and excluded basal resources for primary removal. Robustness was calculated using the methods stated in Dunne and Williams^5^.

### Functional characteristics

We explored functional characteristics of local food-webs shaped by the broad functional feeding groups as described in the previous subsection (see Fig. 1 and Fig. S2). The feeding ecology (i.e., behaviour and diet) that defines each of these groups are further detailed in Supplementary Table S2. To calculate the local functional diversity, we used the number of feeding groups that had at least one genus present in the sample. Functional redundancy was calculated as α-diversity divided by functional diversity, reflecting the average number of genera within a feeding group.

### Data analysis

We ran linear mixed-effects model analysis to assess the relationship between group-specific genus α-diversity (bacteria, invertebrates and fish), food-web structural characteristics (coherence, connectance, link density, number of links, omnivory, modularity, nestedness and robustness) and functional characteristics (functional diversity and redundancy) with site location within the catchment. Drainage area (km^2^) was log transformed to fit model assumptions (normality). For each dependant variable, drainage area and season were the fixed effects. We included site as a random effect to account for repeated sampling of sites. We tested models with and without interactions to determine the significance of this potential interaction (see Supplementary Table S5):

Interactive model: genus α-diversity ~ drainage area * season + (1 | Site)

No Interaction model: genus α-diversity ~ drainage area + season + (1 | Site)

Significance was calculated using the lmerTest R-package^76^, which applies Satterthwaite’s method to estimate degrees of freedom and generate p-values for mixed models (Supplementary Table S6 and S7). We then used anova() to see the overall effect of both drainage area and season (Supplementary Table S8). Subsequent contrast testing were carried out using the emmeans R-package^77^ with the function emtrends() for mixed-effects model slopes contrast testing (Supplementary Table S9), and the function emmeans() for testing significant differences between seasons only (i.e., collapsing the drainage area axis; Supplementary Table S10).

## Acknowledgments

We thank Samuel Hürlemann and Jeanine Brantschen for their help in the laboratory and field. We also thank Xing Xing, Silvana Kaeser, Elvira Mächler and Roman Alther for their assistance with field sample collection, Silvia Kobel and Aria Minder (GDC) for their help with library preparation, and two anonymous reviewers for constructive comments. The data generation and analysis in this paper was generated in collaboration with the Genetic Diversity Centre (https://gdc.ethz.ch), ETH Zurich. Funding is from the University of Zurich Research Priority Programme Global Change and Biodiversity and the Swiss National Science Foundation (grants No. PP00P3_179089 and 31003A_173074) to FA.

## Author contributions

R.C.B and F.A conceived the study. R.C.B performed the genomic lab analyses and J.C.W. performed the bioinformatics. R.C.B assigned trophic levels to all taxa and constructed the metaweb, and H.H performed food-web analysis. R.C.B performed all statistical analyses. R.C.B produced all figures and illustrations. All authors contributed to the interpretation of the networks and discussed and commented on the paper.

## Materials & Correspondence

Correspondence to Rosetta C. Blackman or Florian Altermatt

## Data availability

Sequencing data generated during this study will be made available in a public repository and the data files used in the analysis will be made available upon publication on the following GitHub repository: https://github.com/RosettaBlackman/RiverDNA

