## Supplementary information for "Spatio-temporal patterns of multi-trophic biodiversity and food-web characteristics uncovered across a river catchment using environmental DNA"

*** Corresponding Authors:**

**ORCID IDs:**

Rosetta C. Blackman - https://orcid.org/0000-0002-6182-8691

Hsi-Cheng Ho - https://orcid.org/0000-0002-0734-0249

Jean-Claude Walser - https://orcid.org/0000-0003-1513-0783

Florian Altermatt- https://orcid.org/0000-0002-4831-6958

### Supporting Information

**Figures and Tables:**

Figure S1: Drainage Area
Figure S2: Site genus richness from eDNA samples across all groups and Seasons
Figure S3: Food-web network with horizontal categories.
Figure S4: Supplementary food-web characteristics over space and time (Coherence, Number of links, Modularity, and Robustness)
Figure S5: Local food web structure: visualisation of the local food web at Site G_14

Table S1: MiSeq library loading and output information
Table S2: Function Feeding group categories
Table S3: Food web classifications
Table S4: Alpha diversity from eDNA samples
Table S5: Model comparison for mixed effect models with and without Drainage Area – Season interactions
Table S6: Mixed effect model outputs for all variables tested in this study - fixed effects outputs
Table S7: Mixed model outputs for all variables tested in this study - random effect outputs
Table S8: Fixed effect analysis of variants tables for all interaction models
Table S9: Contrast testing output from emtrends. The pairwise comparison of the mixed effect model slopes.
Table S10: Contrast testing output from emmeans. The pairwise comparison of means between seasons for all variables included in the analysis.
Table S11: β-diversity against river distance for each group and each season.
Table S12: Primer selection for library preparation
Table S13: Positive sample information

**Methods:**
First PCR for library preparation (12S, COI and 16S)
Data preparation workflow - steps and parameters (12S, COI and 16S)

**Figures and Tables:**

**Figure S1: Drainage Area:** Drainage area per site sampled in this study.

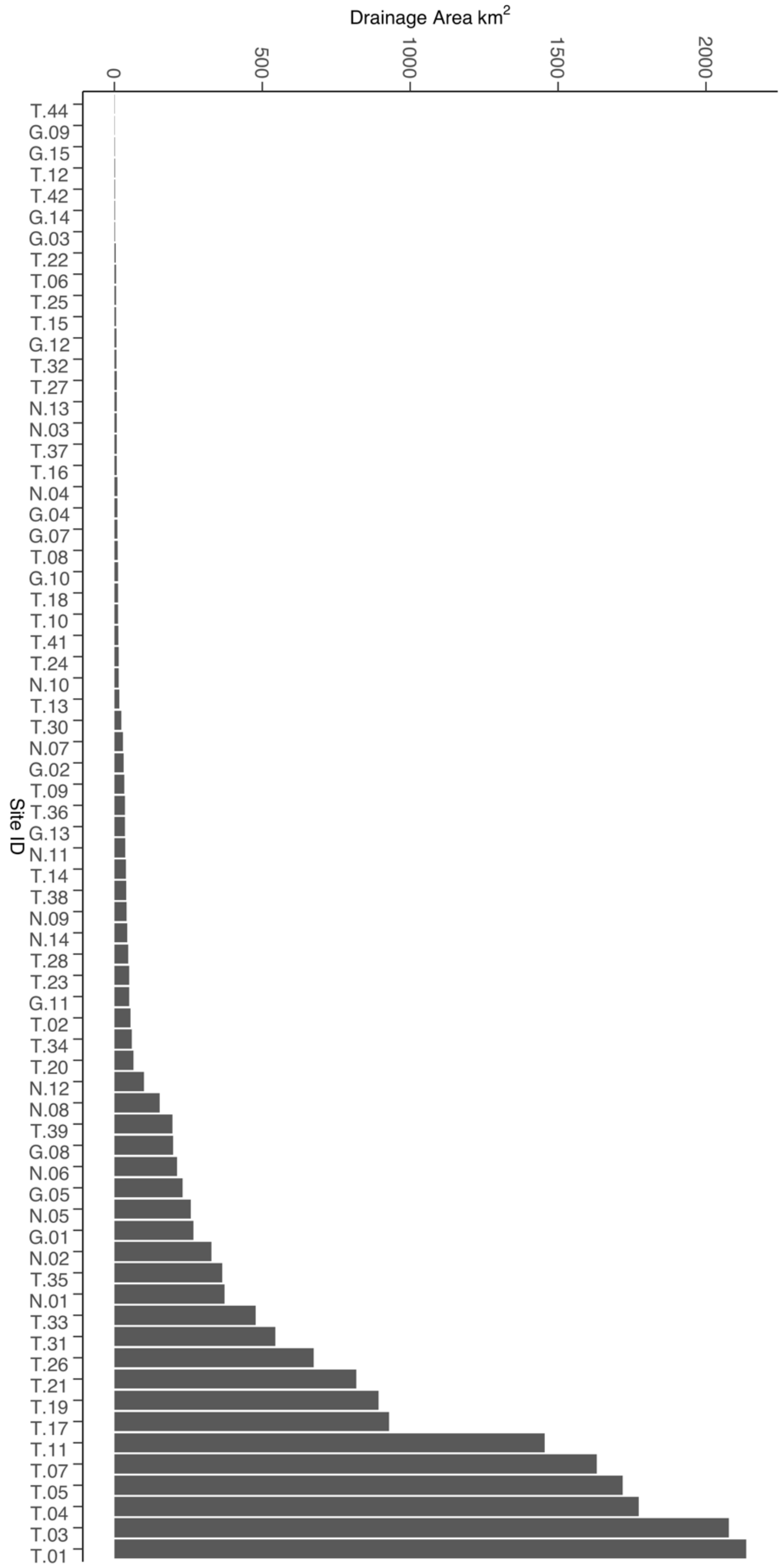

**Figure S2: Site α-diversity (genus richness) from eDNA samples across all groups and Seasons:** Bubble plot to show absolute number of genera recovered in each group from eDNA samples collected in Spring, Summer and Autumn.

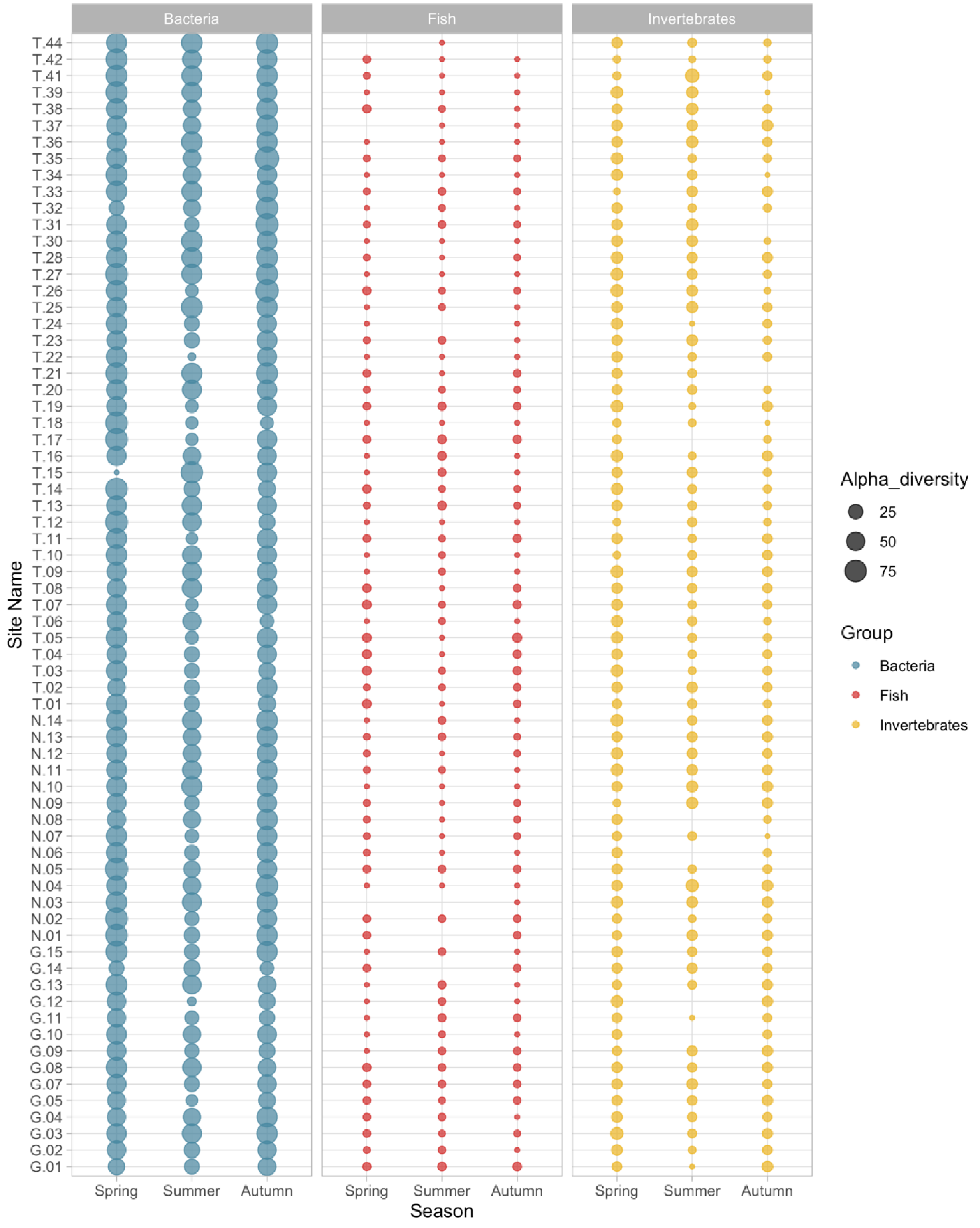

**Figure S3: Food-web network with horizontal categories.** This schematic shows the general groupings of the Functional Feeding Groups (FFG) present in our study: basal resource, decomposer, herbivore/detrivore and predator. This demonstrates the overlap some FFG may have in the overall food web. Colours and labels indicate the FFG.

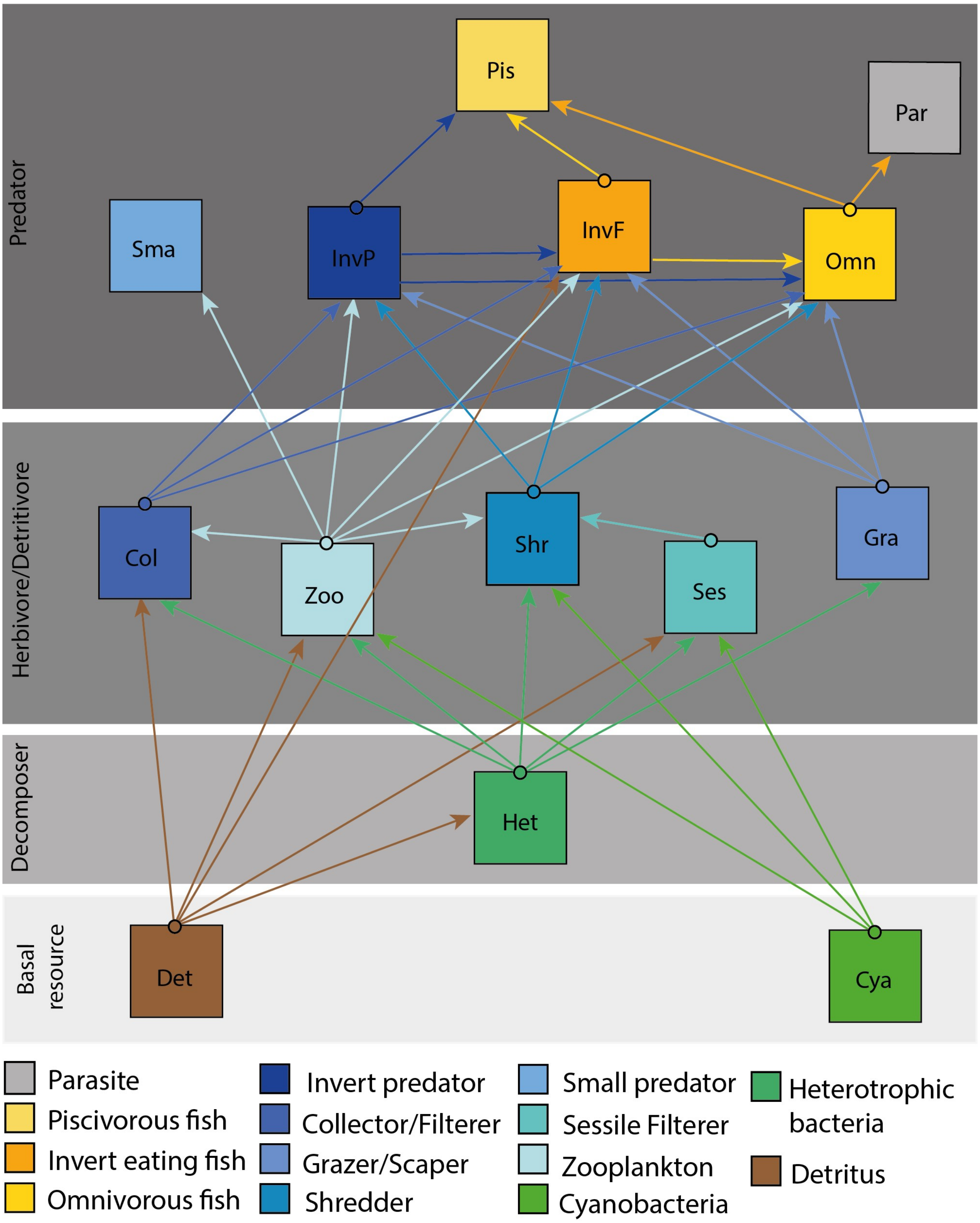

**Figure S4: Supplementary food-web structural characteristics:** Plots show foodweb structural characteristic: a and b: Coherence, c and d: Number of links, e and f: Modularity and g and h: Robustness. Plots a, c, e and g lines indicate linear mixed effect models with shaded area showing 95% confidence intervals as calculated using the model predictions and standard error. Plots a, c and e show the mixed effect models with a significant interaction with the difference across each season (colour represents season). Plot g shows no significant effect of the interaction between drainage area and season; therefore the blue line indicates the mixed effect model output of all the data as a function of drainage area in blue. Plots b, d, f and h show the change in characteristic over season with samples sites linked by grey lines over the three sampling seasons. Colour represents season: yellow – Spring, green – Summer and orange – Autumn.

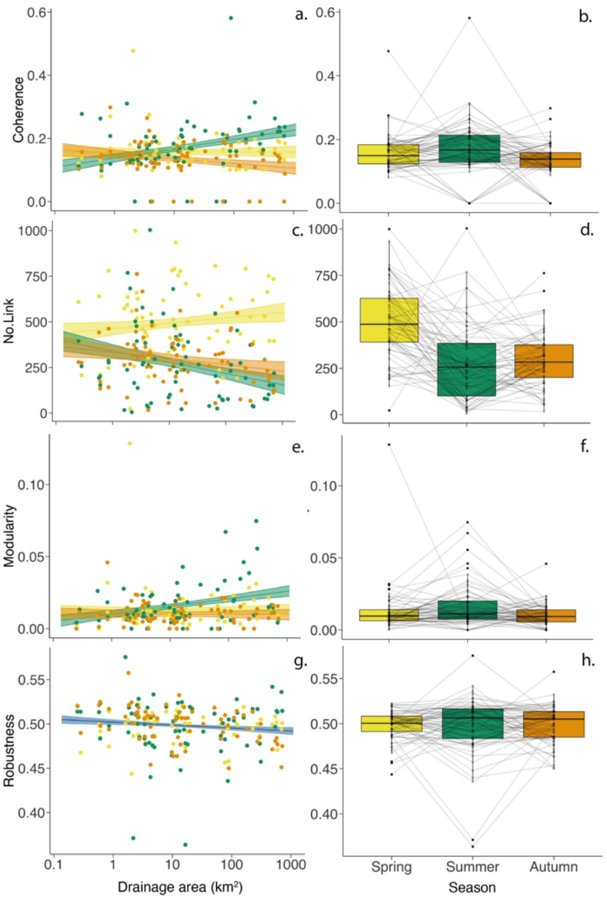

**Figure S5: Local food web structure**: visualisation of the local food web at Site G_14 across Spring, Summer and Autumn as generated using R package Cheddar.

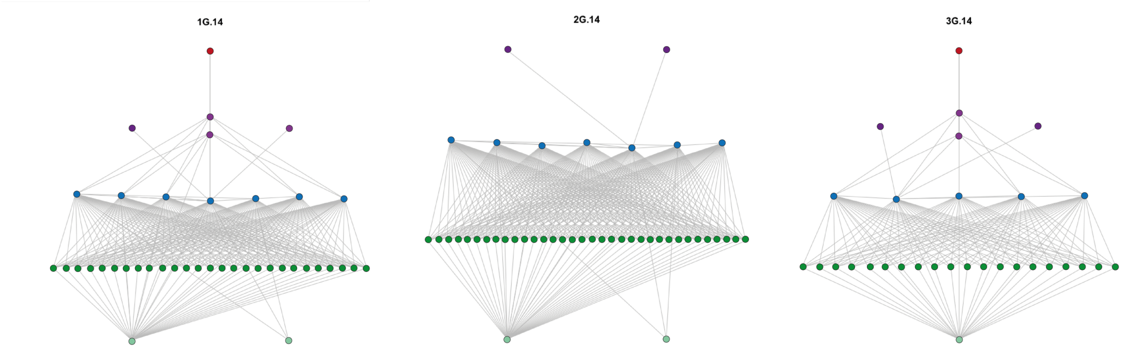

**Table S1: MiSeq library loading, output and bioinformatics information.**

| | **12S** | **COI** | **16S** |
|---|---|---|---|
| **MiSeq regent kit** | 250 cycles PE v2 | 600 cycle PE v3 | 600 cycle PE v3 |
| **Target fragment** | 278 bp | 538 bp | 450 bp |
| **library conc loaded** | 17 pM | 15 pM | 15 pM |
| **PhiX conc loaded** | 10 % | 10 % | 10 % |
| **Read lengths** | 2x 125 cycles | 2x 300 cycles | 2x 300 cycles |
| **Number of raw reads** | 12,291,563 | 14,325,071 | 14,027,834 |
| **Q30** | 95.8% | 84.5% | 77.5% |
| **After data processing** | 10,880,390 | 12,777,525 | 12,563,934 |
| **Average sequencing depth per sample** | 43,176 | 50,704 | 49,857. |
| **OTUs (abundance threshold 2)** | 371 | 13,009 | 34,102 |
| **zOTUs (abundance threshold 7)** | 252 | 6,685 | 30,788 |
| **zOTUs with 97% clustering** | 159 | 3,179 | 11,320 |

**Table S2: Functional Feeding group categories**: Descriptions used to establish interactives between genera of each group:

| **Functional Feeding Group** | **Description** |
|---|---|
| Parasite | Feeds from host |
| Piscivorous fish | Feeds on fish |
| Invertebrate eating fish | Feeds on macroinvertebrates |
| Omnivorous fish | Feeds on both macroinvertebrate and detritus |
| Invertebrate predator | Macroinvertebrate which feeds on macroinvertebrates such as predatory Plecopterans |
| Collector/Filterer | Feeds on sedimented fine particle organic matter |
| Grazer/Scraper | Feeds on bacteria, biofilm, particulate organic matter, and living plants |
| Shredder | Feeds on fallen leaves, plant tissue and coarse organic matter |
| Small Predator | Small predators which feed on microscopic invertebrates and zooplankton |
| Sessile filterer | Non-mobile microscopic filter feeder which feed on bacteria and are grazed on by higher trophic levels (macroinvertebrate) |
| Zooplankton | The animal component of a planktonic community feeding on detritus, bacteria, biofilms and in some case other zooplankton |
| Heterotrophic bacteria | Bacteria which use organic carbon as a resource as opposed to autotrophs |
| Cyanobacteria | Basal resource, uses sunlight to generate resource |
| Detritus | This is the allochthonous inputs which consist of terrestrially derived carbon and forms the basal resource of aquatic food webs. |

**Table S3: Food web classification:** All aquatic associated genus were classified into Functional Feeding groups  to construct the metaweb used for food web analysis. This table has the details of which groups each genus was classified into, a general descriptor of that group and the information source used to derive the groupings.

| Taxa | Library | FFG | General descriptor | Reference |
|---|---|---|---|---|
| **Acanthocyclops** | COI | Zooplankton | Zooplankton | Expert knowledge |
| **Acetobacteroides** | 16S | Heterotroph bac | Bacteria | [1] |
| **Acholeplasma** | 16S | Heterotroph bac | Bacteria | [1] |
| **Achromobacter** | 16S | Heterotroph bac | Bacteria | [1] |
| **Acidibacter** | 16S | Heterotroph bac | Bacteria | [1] |
| **Acidiphilium** | 16S | Heterotroph bac | Bacteria | [1] |
| **Acidovorax** | 16S | Heterotroph bac | Bacteria | [1] |
| **Acinetobacter** | 16S | Heterotroph bac | Bacteria | [1] |
| **Actinoplanes** | 16S | Heterotroph bac | Bacteria | [1] |
| **Aeromonas** | 16S | Heterotroph bac | Bacteria | [1] |
| **Aetherobacter** | 16S | Heterotroph bac | Bacteria | [1] |
| **AKYG587** | 16S | Heterotroph bac | Bacteria | [1] |
| **Algoriphagus** | 16S | Heterotroph bac | Bacteria | [1] |
| **Alistipes** | 16S | Heterotroph bac | Bacteria | [1] |
| **Alkanindiges** | 16S | Heterotroph bac | Bacteria | [1] |
| **Alloprevotella** | 16S | Heterotroph bac | Bacteria | [1] |
| **Altererythrobacter** | 16S | Heterotroph bac | Bacteria | [1] |
| **Amphinemura** | COI | Grazer scraper | Invertebrate | [2] |
| **Anaerocella** | 16S | Heterotroph bac | Bacteria | [1] |
| **Anopheles** | COI | Grazer scraper | Diptera | [3] |
| **Antocha** | COI | Collector Filterer | Diptera | Expert knowledge |
| **Aquabacterium** | 16S | Heterotroph bac | Bacteria | [1] |
| **Aquaspirillum** | 16S | Heterotroph bac | Bacteria | [1] |
| **Aquicella** | 16S | Heterotroph bac | Bacteria | [1] |
| **Aquimonas** | 16S | Heterotroph bac | Bacteria | [1] |
| **Arcicella** | 16S | Heterotroph bac | Bacteria | [1] |
| **Arcobacter** | 16S | Heterotroph bac | Bacteria | [1] |
| **Arenimonas** | 16S | Heterotroph bac | Bacteria | [1] |
| **Armatimonas** | 16S | Heterotroph bac | Bacteria | [1] |
| **Asaia** | 16S | Heterotroph bac | Bacteria | [1] |
| **Asellus** | COI | Collector Filterer | Invertebrate | [4] |
| **Asticcacaulis** | 16S | Heterotroph bac | Bacteria | [1] |
| **Azospirillum** | 16S | Heterotroph bac | Bacteria | [1] |

| | | | | |
|---|---|---|---|---|
| **Bacillus** | 16S | Heterotroph bac | Bacteria | [1] |
| **Bacteriovorax** | 16S | Heterotroph bac | Bacteria | [1] |
| **Bacteroides** | 16S | Heterotroph bac | Bacteria | [1] |
| **Baetis** | COI | Grazer scraper | Invertebrate | [5] |
| **Barbatula** | COI | Invert eating fish | Fish | [6] |
| **Barbus** | 12S | Invert eating fish | Fish | [6] |
| **Bauldia** | 16S | Heterotroph bac | Bacteria | [1] |
| **Bdellovibrio** | 16S | Heterotroph bac | Bacteria | [1] |
| **Beggiatoa** | 16S | Heterotroph bac | Bacteria | [1] |
| **Bifidobacterium** | 16S | Heterotroph bac | Bacteria | [1] |
| **Blastocatella** | 16S | Heterotroph bac | Bacteria | [1] |
| **Blautia** | 16S | Heterotroph bac | Bacteria | [1] |
| **Bosea** | 16S | Heterotroph bac | Bacteria | [1] |
| **Brevundimonas** | 16S | Heterotroph bac | Bacteria | [1] |
| **Bryobacter** | 16S | Heterotroph bac | Bacteria | [1] |
| **BSV13** | 16S | Heterotroph bac | Bacteria | [1] |
| **Caenis** | COI | Collector Filterer | Invertebrate | [7] |
| **Capnioneura** | COI | Shredder | Invertebrate | [8] |
| **Carassius** | COI | Omnivorous fish | Fish | [6] |
| **Catenibacterium** | 16S | Heterotroph bac | Bacteria | [1] |
| **Caulobacter** | 16S | Heterotroph bac | Bacteria | [1] |
| **Cellvibrio** | 16S | Heterotroph bac | Bacteria | [1] |
| **Chaetogaster** | COI | small preds | Invertebrate | [9] |
| **Chaetonotus** | COI | Zooplankton | Gastrotrich | Expert knowledge |
| **Chamaesiphon** | 16S | Autotroph bac | Bacteria | [1] |
| **Chironomus** | COI | Collector Filterer | Diptera | [10] |
| **Chitinibacter** | 16S | Heterotroph bac | Bacteria | [1] |
| **Chitinimonas** | 16S | Heterotroph bac | Bacteria | [1] |
| **Chitinivorax** | 16S | Heterotroph bac | Bacteria | [1] |
| **Chitinophaga** | 16S | Heterotroph bac | Bacteria | [1] |
| **Chroococcidiopsis** | 16S | Heterotroph bac | Bacteria | [1] |
| **Chryseobacterium** | 16S | Heterotroph bac | Bacteria | [1] |
| **Chryseolinea** | 16S | Heterotroph bac | Bacteria | [1] |
| **Chthoniobacter** | 16S | Heterotroph bac | Bacteria | [1] |
| **Chydorus** | COI | Zooplankton | Zooplankton | Expert knowledge |
| **Cloacibacterium** | 16S | Heterotroph bac | Bacteria | [1] |
| **Cloeon** | COI | Grazer scraper | Invertebrate | [5] |
| **Clostridium** | 16S | Heterotroph bac | Bacteria | [1] |
| **Collinsella** | 16S | Heterotroph bac | Bacteria | [1] |

| | | | | |
|---|---|---|---|---|
| **Comamonas** | 16S | Heterotroph bac | Bacteria | [1] |
| **Conchapelopia** | COI | Invert predator | Diptera | [10] |
| **Corynebacterium 1** | 16S | Heterotroph bac | Bacteria | [1] |
| **Cottus** | 12S | Invert eating fish | Fish | [6] |
| **Coxiella** | 16S | Heterotroph bac | Bacteria | [1] |
| **Craspedacusta** | COI | small preds | Hydrozoan | Expert knowledge |
| **Crenothrix** | 16S | Heterotroph bac | Bacteria | [1] |
| **Cricotopus** | COI | Grazer scraper | Diptera | [10] |
| **Crocinitomix** | 16S | Heterotroph bac | Bacteria | [1] |
| **Cyclops** | COI | Zooplankton | Zooplankton | Expert knowledge |
| **Cypridopsis** | COI | Zooplankton | Zooplankton | Expert knowledge |
| **Cyprinus** | COI | Invert eating fish | Fish | Expert knowledge |
| **Cytophaga** | 16S | Heterotroph bac | Bacteria | [1] |
| **Deefgea** | 16S | Heterotroph bac | Bacteria | [1] |
| **Deinococcus** | 16S | Heterotroph bac | Bacteria | [1] |
| **Delftia** | 16S | Heterotroph bac | Bacteria | [1] |
| **Dero** | COI | Collector Filterer | Invertebrate | Expert knowledge |
| **Detritus** | NA | resource | leaf litter | Expert knowledge |
| **Devosia** | 16S | Heterotroph bac | Bacteria | [1] |
| **Diamesa** | COI | Grazer scraper | Diptera | [10] |
| **Dickeya** | 16S | Heterotroph bac | Bacteria | [1] |
| **Dicranomyia** | COI | Shredder | Diptera | Expert knowledge |
| **Dietzia** | 16S | Heterotroph bac | Bacteria | [1] |
| **Dinghuibacter** | 16S | Heterotroph bac | Bacteria | [1] |
| **Dongia** | 16S | Heterotroph bac | Bacteria | [1] |
| **Duganella** | 16S | Heterotroph bac | Bacteria | [1] |
| **Dyadobacter** | 16S | Heterotroph bac | Bacteria | [1] |
| **Dysgonomonas** | 16S | Heterotroph bac | Bacteria | [1] |
| **Ecdyonurus** | COI | Grazer scraper | Invertebrate | [5] |
| **Eiseniella** | COI | Collector Filterer | Invertebrate | [9] |
| **Elmis** | COI | Grazer scraper | Invertebrate | [11] |
| **Elodes** | COI | Shredder | Invertebrate | Expert knowledge |
| **Elstera** | 16S | Heterotroph bac | Bacteria | [1] |
| **Emticicia** | 16S | Heterotroph bac | Bacteria | [1] |
| **Enhydrobacter** | 16S | Heterotroph bac | Bacteria | [1] |
| **Epeorus** | COI | Grazer scraper | Invertebrate | [5] |
| **Ephemera** | COI | Collector Filterer | Invertebrate | [5] |
| **Ephemerella** | COI | Grazer scraper | Invertebrate | [5] |
| **Ephydatia** | COI | sessile filterers | Sponge | Expert knowledge |

| | | | | |
|---|---|---|---|---|
| **Erysipelothrix** | 16S | Heterotroph bac | Bacteria | [1] |
| **Esox** | 12S | Piscivore fish | Fish | [6] |
| **Euchlanis** | COI | Zooplankton | Zooplankton | Expert knowledge |
| **Eucyclops** | COI | Zooplankton | Zooplankton | Expert knowledge |
| **Euglenaria anabaena** | 16S | Heterotroph bac | Bacteria | [1] |
| **Eukiefferiella** | COI | Grazer scraper | Diptera | [10] |
| **Exiguobacterium** | 16S | Heterotroph bac | Bacteria | [1] |
| **Facklamia** | 16S | Heterotroph bac | Bacteria | [1] |
| **Faecalibacterium** | 16S | Heterotroph bac | Bacteria | [1] |
| **Fastidiosipila** | 16S | Heterotroph bac | Bacteria | [1] |
| **Ferruginibacter** | 16S | Heterotroph bac | Bacteria | [1] |
| **Fibrella** | 16S | Heterotroph bac | Bacteria | [1] |
| **Filimonas** | 16S | Heterotroph bac | Bacteria | [1] |
| **Flavihumibacter** | 16S | Heterotroph bac | Bacteria | [1] |
| **Flavitalea** | 16S | Heterotroph bac | Bacteria | [1] |
| **Flavobacterium** | 16S | Heterotroph bac | Bacteria | [1] |
| **Flectobacillus** | 16S | Heterotroph bac | Bacteria | [1] |
| **Fluviicoccus** | 16S | Heterotroph bac | Bacteria | [1] |
| **Fluviicola** | 16S | Heterotroph bac | Bacteria | [1] |
| **Fluviimonas** | 16S | Heterotroph bac | Bacteria | [1] |
| **Fodinicola** | 16S | Heterotroph bac | Bacteria | [1] |
| **Fusibacter** | 16S | Heterotroph bac | Bacteria | [1] |
| **Fusobacterium** | 16S | Heterotroph bac | Bacteria | [1] |
| **Gaiella** | 16S | Heterotroph bac | Bacteria | [1] |
| **Gammarus** | COI | Shredder | Invertebrate | [4] |
| **Gemmata** | 16S | Heterotroph bac | Bacteria | [1] |
| **Gemmatimonas** | 16S | Heterotroph bac | Bacteria | [1] |
| **Geobacter** | 16S | Heterotroph bac | Bacteria | [1] |
| **GKS98 freshwater group** | 16S | Heterotroph bac | Bacteria | [1] |
| **Glossosoma** | COI | Grazer scraper | Invertebrate | Expert knowledge |
| **Gobio** | 12S | Invert eating fish | Fish | [6] |
| **Granulicella** | 16S | Heterotroph bac | Bacteria | [1] |
| **Gyrodactylus** | COI | Parasite | Parasite | Expert knowledge |
| **H16** | 16S | Heterotroph bac | Bacteria | [1] |
| **Habroleptoides** | COI | Collector Filterer | Invertebrate | [5] |
| **Habrophlebia** | COI | Collector Filterer | Invertebrate | [5] |
| **Haematospirillum** | 16S | Heterotroph bac | Bacteria | [1] |
| **Haliangium** | 16S | Heterotroph bac | Bacteria | [1] |
| **Haliea** | 16S | Heterotroph bac | Bacteria | [1] |

| | | | | |
|---|---|---|---|---|
| **Halioglobus** | 16S | Heterotroph bac | Bacteria | [1] |
| **Haliscomenobacter** | 16S | Heterotroph bac | Bacteria | [1] |
| **Haloferula** | 16S | Heterotroph bac | Bacteria | [1] |
| **Halomonas** | 16S | Heterotroph bac | Bacteria | [1] |
| **Herpetosiphon** | 16S | Heterotroph bac | Bacteria | [1] |
| **hgcI clade** | 16S | Heterotroph bac | Bacteria | [1] |
| **Hirschia** | 16S | Heterotroph bac | Bacteria | [1] |
| **Holdemanella** | 16S | Heterotroph bac | Bacteria | [1] |
| **Hydra** | COI | small preds | Hydrozoan | Expert knowledge |
| **Hydrogenophaga** | 16S | Heterotroph bac | Bacteria | [1] |
| **Hydropsyche** | COI | Collector Filterer | Invertebrate | [8] |
| **Hydroptila** | COI | Collector Filterer | Invertebrate | [8] |
| **Hymenobacter** | 16S | Heterotroph bac | Bacteria | [1] |
| **Hyphomicrobium** | 16S | Heterotroph bac | Bacteria | [1] |
| **Iamia** | 16S | Heterotroph bac | Bacteria | [1] |
| **Ilumatobacter** | 16S | Heterotroph bac | Bacteria | [1] |
| **Iodobacter** | 16S | Heterotroph bac | Bacteria | [1] |
| **Isohypsibius** | COI | Zooplankton | Invertebrate | Expert knowledge |
| **Isoperla** | COI | Invert predator | Invertebrate | [8] |
| **Keratella** | COI | Zooplankton | Zooplankton | Expert knowledge |
| **Kurthia** | 16S | Heterotroph bac | Bacteria | [1] |
| **Lachnoclostridium** | 16S | Heterotroph bac | Bacteria | [1] |
| **Lacibacter** | 16S | Heterotroph bac | Bacteria | [1] |
| **Lactobacillus** | 16S | Heterotroph bac | Bacteria | [1] |
| **Lactococcus** | 16S | Heterotroph bac | Bacteria | [1] |
| **Leadbetterella** | 16S | Heterotroph bac | Bacteria | [1] |
| **Leeia** | 16S | Heterotroph bac | Bacteria | [1] |
| **Legionella** | 16S | Heterotroph bac | Bacteria | [1] |
| **Leptolyngbya** | 16S | Autotroph bac | Bacteria | [1] |
| **Leptothrix** | 16S | Heterotroph bac | Bacteria | [1] |
| **Leptotrichia** | 16S | Heterotroph bac | Bacteria | [1] |
| **Leucobacter** | 16S | Heterotroph bac | Bacteria | [1] |
| **Leuctra** | COI | Collector Filterer | Invertebrate | [8] |
| **Limnephilus** | COI | Shredder | Invertebrate | [8] |
| **Limnius** | COI | Grazer scraper | Invertebrate | [11] |
| **Limnobacter** | 16S | Heterotroph bac | Bacteria | [1] |
| **Limnodrilus** | COI | Collector Filterer | Invertebrate | [9] |
| **Limnohabitans** | 16S | Heterotroph bac | Bacteria | [1] |
| **Limnophyes** | COI | Collector Filterer | Diptera | Expert knowledge |

| | | | | |
|---|---|---|---|---|
| **Lumbricillus** | COI | Collector Filterer | Invertebrate | [9] |
| **Luteimonas** | 16S | Heterotroph bac | Bacteria | [1] |
| **Luteolibacter** | 16S | Heterotroph bac | Bacteria | [1] |
| **Lysobacter** | 16S | Heterotroph bac | Bacteria | [1] |
| **Macellibacteroides** | 16S | Heterotroph bac | Bacteria | [1] |
| **Magnetospirillum** | 16S | Heterotroph bac | Bacteria | [1] |
| **Marinospirillum** | 16S | Heterotroph bac | Bacteria | [1] |
| **Massilia** | 16S | Heterotroph bac | Bacteria | [1] |
| **Megamonas** | 16S | Heterotroph bac | Bacteria | [1] |
| **Meganema** | 16S | Heterotroph bac | Bacteria | [1] |
| **Melampophylax** | COI | Shredder | Invertebrate | [8] |
| **Merismopedia** | 16S | Heterotroph bac | Bacteria | [1] |
| **Mesorhizobium** | 16S | Heterotroph bac | Bacteria | [1] |
| **Methanogenium** | 16S | Heterotroph bac | Bacteria | [1] |
| **Methylobacterium** | 16S | Heterotroph bac | Bacteria | [1] |
| **Methyloparacoccus** | 16S | Heterotroph bac | Bacteria | [1] |
| **Methylophilus** | 16S | Heterotroph bac | Bacteria | [1] |
| **Methylotenera** | 16S | Heterotroph bac | Bacteria | [1] |
| **Microbacterium** | 16S | Heterotroph bac | Bacteria | [1] |
| **Microcoleus** | 16S | Heterotroph bac | Bacteria | [1] |
| **Micropsectra** | COI | Collector Filterer | Diptera | [10] |
| **Mucilaginibacter** | 16S | Heterotroph bac | Bacteria | [1] |
| **Mycobacterium** | 16S | Heterotroph bac | Bacteria | [1] |
| **Nakamurella** | 16S | Heterotroph bac | Bacteria | [1] |
| **Nannocystis** | 16S | Heterotroph bac | Bacteria | [1] |
| **Nemoura** | COI | Shredder | Invertebrate | [8] |
| **Nevskia** | 16S | Heterotroph bac | Bacteria | [1] |
| **Nitrospira** | 16S | Autotroph bac | Bacteria | [1] |
| **Niveispirillum** | 16S | Heterotroph bac | Bacteria | [1] |
| **Nocardioides** | 16S | Heterotroph bac | Bacteria | [1] |
| **Novosphingobium** | 16S | Heterotroph bac | Bacteria | [1] |
| **Oleiphilus** | 16S | Heterotroph bac | Bacteria | [1] |
| **Oligoflexus** | 16S | Heterotroph bac | Bacteria | [1] |
| **Oligosphaera** | 16S | Heterotroph bac | Bacteria | [1] |
| **OM27 clade** | 16S | Heterotroph bac | Bacteria | [1] |
| **Ophidonais** | COI | Collector Filterer | Invertebrate | [9] |
| **Opitutus** | 16S | Heterotroph bac | Bacteria | [1] |
| **Orthocladius** | COI | Collector Filterer | Diptera | [10] |
| **Paenibacillus** | 16S | Heterotroph bac | Bacteria | [1] |

| | | | | |
|---|---|---|---|---|
| **Paludibacter** | 16S | Heterotroph bac | Bacteria | [1] |
| **Paludibaculum** | 16S | Heterotroph bac | Bacteria | [1] |
| **Parabacteroides** | 16S | Heterotroph bac | Bacteria | [1] |
| **Parafilimonas** | 16S | Heterotroph bac | Bacteria | [1] |
| **Parasediminibacterium** | 16S | Heterotroph bac | Bacteria | [1] |
| **Parasegetibacter** | 16S | Heterotroph bac | Bacteria | [1] |
| **Paratrichocladius** | COI | Grazer scraper | Diptera | [10] |
| **Pedobacter** | 16S | Heterotroph bac | Bacteria | [1] |
| **Pedomicrobium** | 16S | Heterotroph bac | Bacteria | [1] |
| **Pelosinus** | 16S | Heterotroph bac | Bacteria | [1] |
| **Perca** | 12S | Omnivorous fish | Fish | [6] |
| **Peredibacter** | 16S | Heterotroph bac | Bacteria | [1] |
| **Perla** | COI | Invert predator | Invertebrate | [8] |
| **Perlodes** | COI | Invert predator | Invertebrate | [8] |
| **Perlucidibaca** | 16S | Heterotroph bac | Bacteria | [1] |
| **Petrimonas** | 16S | Heterotroph bac | Bacteria | [1] |
| **Phacotus lenticularis** | 16S | Heterotroph bac | Bacteria | [1] |
| **Phaeodactylibacter** | 16S | Heterotroph bac | Bacteria | [1] |
| **Phascolarctobacterium** | 16S | Heterotroph bac | Bacteria | [1] |
| **Phaselicystis** | 16S | Heterotroph bac | Bacteria | [1] |
| **Phenylobacterium** | 16S | Heterotroph bac | Bacteria | [1] |
| **Philopotamus** | COI | Collector Filterer | Invertebrate | [8] |
| **Phormidium** | 16S | Heterotroph bac | Bacteria | [1] |
| **Phoxinus** | 12S | Invert eating fish | Fish | [6] |
| **Phycisphaera** | 16S | Heterotroph bac | Bacteria | [1] |
| **Pir4 lineage** | 16S | Heterotroph bac | Bacteria | [1] |
| **Pirellula** | 16S | Heterotroph bac | Bacteria | [1] |
| **Planctomyces** | 16S | Heterotroph bac | Bacteria | [1] |
| **Planktothrix** | 16S | Autotroph bac | Bacteria | [1] |
| **Pleurocapsa** | 16S | Heterotroph bac | Bacteria | [1] |
| **Plumatella** | COI | sessile filterers | Bryozoan | [12] |
| **Polaromonas** | 16S | Heterotroph bac | Bacteria | [1] |
| **Polyangium** | 16S | Heterotroph bac | Bacteria | [1] |
| **Polyarthra** | COI | Zooplankton | Zooplankton | Expert knowledge |
| **Polymorphobacter** | 16S | Heterotroph bac | Bacteria | [1] |
| **Polynucleobacter** | 16S | Heterotroph bac | Bacteria | [1] |
| **Polypedilum** | COI | Collector Filterer | Diptera | [10] |
| **Potamophylax** | COI | Shredder | Invertebrate | [8] |
| **Potamopyrgus** | COI | Collector Filterer | Invertebrate | [13] |

| | | | | |
|---|---|---|---|---|
| **Potamothrix** | COI | Collector Filterer | Invertebrate | [9] |
| **Prevotella 9** | 16S | Heterotroph bac | Bacteria | [1] |
| **Pristina** | COI | Collector Filterer | Invertebrate | [9] |
| **Prodiamesa** | COI | Collector Filterer | Diptera | [10] |
| **Propionivibrio** | 16S | Heterotroph bac | Bacteria | [1] |
| **Prosimulium** | COI | Collector Filterer | Diptera | [3] |
| **Prosthecobacter** | 16S | Heterotroph bac | Bacteria | [1] |
| **Proteiniclasticum** | 16S | Heterotroph bac | Bacteria | [1] |
| **Proteiniphilum** | 16S | Heterotroph bac | Bacteria | [1] |
| **Protonemura** | COI | Shredder | Invertebrate | [8] |
| **Pseudarcicella** | 16S | Heterotroph bac | Bacteria | [1] |
| **Pseudendoclonium akinetum** | 16S | Heterotroph bac | Bacteria | [1] |
| **Pseudohongiella** | 16S | Heterotroph bac | Bacteria | [1] |
| **Pseudomonas** | 16S | Heterotroph bac | Bacteria | [1] |
| **Pseudorhodoferax** | 16S | Heterotroph bac | Bacteria | [1] |
| **Pseudoxanthomonas** | 16S | Heterotroph bac | Bacteria | [1] |
| **Psychrobacter** | 16S | Heterotroph bac | Bacteria | [1] |
| **Reyranella** | 16S | Heterotroph bac | Bacteria | [1] |
| **Rheinheimera** | 16S | Heterotroph bac | Bacteria | [1] |
| **Rhithrogena** | COI | Grazer scraper | Invertebrate | [5] |
| **Rhizobacter** | 16S | Heterotroph bac | Bacteria | [1] |
| **Rhizobium** | 16S | Heterotroph bac | Bacteria | [1] |
| **Rhizomicrobium** | 16S | Heterotroph bac | Bacteria | [1] |
| **Rhizorhapis** | 16S | Heterotroph bac | Bacteria | [1] |
| **Rhodobacter** | 16S | Heterotroph bac | Bacteria | [1] |
| **Rhodococcus** | 16S | Heterotroph bac | Bacteria | [1] |
| **Rhodoferax** | 16S | Heterotroph bac | Bacteria | [1] |
| **Rhodomicrobium** | 16S | Heterotroph bac | Bacteria | [1] |
| **Rhyacophila** | COI | Invert predator | Invertebrate | [8] |
| **Rickettsia** | 16S | Heterotroph bac | Bacteria | [1] |
| **Rickettsiella** | 16S | Heterotroph bac | Bacteria | [1] |
| **Riolus** | COI | Grazer scraper | Invertebrate | [11] |
| **Roseiflexus** | 16S | Autotroph bac | Bacteria | [1] |
| **Roseococcus** | 16S | Heterotroph bac | Bacteria | [1] |
| **Roseomonas** | 16S | Heterotroph bac | Bacteria | [1] |
| **Rubellimicrobium** | 16S | Heterotroph bac | Bacteria | [1] |
| **Rudanella** | 16S | Heterotroph bac | Bacteria | [1] |
| **Runella** | 16S | Heterotroph bac | Bacteria | [1] |
| **Salmo** | COI | Invert eating fish | Fish | [6] |

| | | | | |
|---|---|---|---|---|
| **Sandaracinobacter** | 16S | Heterotroph bac | Bacteria | [1] |
| **Sandaracinus** | 16S | Heterotroph bac | Bacteria | [1] |
| **Sandarakinorhabdus** | 16S | Heterotroph bac | Bacteria | [1] |
| **Sarcina** | 16S | Heterotroph bac | Bacteria | [1] |
| **Scardinius** | COI | Omnivorous fish | Fish | [6] |
| **Schlesneria** | 16S | Heterotroph bac | Bacteria | [1] |
| **Sediminibacterium** | 16S | Heterotroph bac | Bacteria | [1] |
| **Sericostoma** | COI | Shredder | Invertebrate | [8] |
| **Serratella** | COI | Grazer scraper | Invertebrate | [5] |
| **Serratia** | 16S | Heterotroph bac | Bacteria | [1] |
| **Sideroxydans** | 16S | Heterotroph bac | Bacteria | [1] |
| **Silanimonas** | 16S | Heterotroph bac | Bacteria | [1] |
| **Simiduia** | 16S | Heterotroph bac | Bacteria | [1] |
| **Simplicispira** | 16S | Heterotroph bac | Bacteria | [1] |
| **Simulium** | COI | Collector Filterer | Diptera | [3] |
| **SM1A02** | 16S | Heterotroph bac | Bacteria | [1] |
| **Snowella** | 16S | Heterotroph bac | Bacteria | [1] |
| **Solitalea** | 16S | Heterotroph bac | Bacteria | [1] |
| **Sorangium** | 16S | Heterotroph bac | Bacteria | [1] |
| **Sphaerochaeta** | 16S | Heterotroph bac | Bacteria | [1] |
| **Sphaerotilus** | 16S | Heterotroph bac | Bacteria | [1] |
| **Sphingobacterium** | 16S | Heterotroph bac | Bacteria | [1] |
| **Sphingobium** | 16S | Heterotroph bac | Bacteria | [1] |
| **Sphingomonas** | 16S | Heterotroph bac | Bacteria | [1] |
| **Sphingopyxis** | 16S | Heterotroph bac | Bacteria | [1] |
| **Sphingorhabdus** | 16S | Heterotroph bac | Bacteria | [1] |
| **Spirochaeta 2** | 16S | Heterotroph bac | Bacteria | [1] |
| **Spirosoma** | 16S | Heterotroph bac | Bacteria | [1] |
| **Sporichthya** | 16S | Heterotroph bac | Bacteria | [1] |
| **Squalius** | COI | Invert eating fish | Fish | [6] |
| **Staphylococcus** | 16S | Heterotroph bac | Bacteria | [1] |
| **Stenostomum** | COI | small preds | Parasite | Expert knowledge |
| **Stenotrophobacter** | 16S | Heterotroph bac | Bacteria | [1] |
| **Stenotrophomonas** | 16S | Heterotroph bac | Bacteria | [1] |
| **Steroidobacter** | 16S | Heterotroph bac | Bacteria | [1] |
| **Streptococcus** | 16S | Heterotroph bac | Bacteria | [1] |
| **Streptomyces** | 16S | Heterotroph bac | Bacteria | [1] |
| **Stylodrilus** | COI | Collector Filterer | Invertebrate | [9] |
| **Sulfurifustis** | 16S | Heterotroph bac | Bacteria | [1] |

| | | | | |
|---|---|---|---|---|
| **Sulfuritalea** | 16S | Heterotroph bac | Bacteria | [1] |
| **Sulfurospirillum** | 16S | Heterotroph bac | Bacteria | [1] |
| **Sutterella** | 16S | Heterotroph bac | Bacteria | [1] |
| **Synechococcus** | 16S | Autotroph bac | Bacteria | [1] |
| **Taibaiella** | 16S | Heterotroph bac | Bacteria | [1] |
| **Takobia** | COI | Grazer scraper | Invertebrate | Expert knowledge |
| **Tanytarsus** | COI | Collector Filterer | Diptera | [10] |
| **Terrimicrobium** | 16S | Heterotroph bac | Bacteria | [1] |
| **Terrimonas** | 16S | Heterotroph bac | Bacteria | [1] |
| **Thauera** | 16S | Heterotroph bac | Bacteria | [1] |
| **Thermocyclops** | COI | Zooplankton | Zooplankton | Expert knowledge |
| **Thermomonas** | 16S | Heterotroph bac | Bacteria | [1] |
| **Thiothrix** | 16S | Heterotroph bac | Bacteria | [1] |
| **Tissierella** | 16S | Heterotroph bac | Bacteria | [1] |
| **Treponema 2** | 16S | Heterotroph bac | Bacteria | [1] |
| **Trichococcus** | 16S | Heterotroph bac | Bacteria | [1] |
| **Truepera** | 16S | Heterotroph bac | Bacteria | [1] |
| **Turicibacter** | 16S | Heterotroph bac | Bacteria | [1] |
| **Tvetenia** | COI | Grazer scraper | Invertebrate | [10] |
| **Uliginosibacterium** | 16S | Heterotroph bac | Bacteria | [1] |
| **Undibacterium** | 16S | Heterotroph bac | Bacteria | [1] |
| **Variibacter** | 16S | Heterotroph bac | Bacteria | [1] |
| **Verrucomicrobium** | 16S | Heterotroph bac | Bacteria | [1] |
| **Vogesella** | 16S | Heterotroph bac | Bacteria | [1] |
| **Woodsholea** | 16S | Heterotroph bac | Bacteria | [1] |
| **Yersinia** | 16S | Heterotroph bac | Bacteria | [1] |
| **Z20** | 16S | Heterotroph bac | Bacteria | [1] |
| **Zoogloea** | 16S | Heterotroph bac | Bacteria | [1] |
| **Zymomonas** | 16S | Heterotroph bac | Bacteria | [1] |
| **12up** | 16S | Heterotroph bac | Bacteria | [1] |

**Table S4: Alpha diversity (Genus richness) from eDNA samples:** reported for each group separately and combined and across each season and combined.

| Group | | Fish | Invertebrates | Bacteria | All genera |
|---|---|---|---|---|---|
| **Spring** | mean α-diversity | 2.068 | 8.849 | 59.384 | 70 |
| | standard deviation | 1.316 | 2.970 | 12.572 | 12.582 |
| | range | 0-6 | 2-15 | 1-81 | 10-92 |
| **Summer** | mean α-diversity | 2.014 | 6.637 | 39.159 | 47.811 |
| | standard deviation | 1.254 | 3.808 | 16.787 | 18.339 |
| | range | 0-5 | 0-20 | 3-73 | 8-85 |
| **Autumn** | mean α-diversity | 1.849 | 5.410 | 55.520 | 62.780 |
| | standard deviation | 1.221 | 2.748 | 14.319 | 13.962 |
| | range | 0-6 | 0-10 | 17-90 | 19-96 |
| **All seasons combined** | mean α-diversity | 1.976 | 6.972 | 51.581 | 60.530 |
| | standard deviation | 1.77 | 3.490 | 16.95 | 17.668 |
| | range | 0-6 | 0-20 | 1-90 | 8-96 |

**Table S5: Model comparison**: mixed effect models with and without Drainage Area – Season interactions. P values < 0.05 indicate cases where the model with an interaction is the significantly better model.

| | Model | npar | AIC | BIC | logLik | deviance | Chisq | Df | Pr(>Chisq) | |
|---|---|---|---|---|---|---|---|---|---|---|
| **Bacteria diversity** | No interaction | 6 | 1702.4 | 1722.4 | -845.2 | 1690.4 | | | | |
| | Interaction | 8 | 1689 | 1715.7 | -836.51 | 1673 | 17.389 | 2 | 0.0001675 | *** |
| **Fish diversity** | No interaction | 6 | 635.97 | 655.97 | -311.99 | 623.97 | | | | |
| | Interaction | 8 | 616.37 | 643.03 | -300.19 | 600.37 | 23.602 | 2 | 7.50E-06 | *** |
| **Invert diversity** | No interaction | 6 | 1069.1 | 1089.1 | -528.55 | 1057.1 | | | | |
| | Interaction | 8 | 1070.6 | 1097.2 | -527.29 | 1054.6 | 2.5147 | 2 | 0.2844 | |
| **No. Link** | No interaction | 6 | 2748.9 | 2768.9 | -1368.4 | 2736.9 | | | | |
| | Interaction | 8 | 2743.8 | 2770.4 | -1363.9 | 2727.8 | 9.0981 | 2 | 0.01058 | * |
| **Link density** | No interaction | 6 | 908.82 | 928.82 | -448.41 | 896.82 | | | | |
| | Interaction | 8 | 908.74 | 935.4 | -446.37 | 892.74 | 4.0792 | 2 | 0.1301 | |
| **Nestedness** | No interaction | 6 | 1169.2 | 1189.1 | -578.57 | 1157.2 | | | | |
| | Interaction | 8 | 1158.1 | 1184.8 | -571.06 | 1142.1 | 15.036 | 2 | 0.0005433 | *** |
| **Coherence** | No interaction | 6 | -514.15 | -494.16 | 263.08 | -526.15 | | | | |
| | Interaction | 8 | -523.68 | -497.02 | 269.84 | -539.68 | 13.528 | 2 | 0.001154 | ** |
| **Connectance** | No interaction | 6 | -784.9 | -764.9 | 398.45 | -796.9 | | | | |
| | Interaction | 8 | -789.76 | -763.09 | 402.88 | -805.76 | 8.8558 | 2 | 0.01194 | * |
| **Omnivory** | No interaction | 6 | -848.11 | -828.11 | 430.05 | -860.11 | | | | |
| | Interaction | 8 | -849.72 | -823.06 | 432.86 | -865.72 | 5.6119 | 2 | 0.06045 | |
| **Robustness** | No interaction | 6 | -952.88 | -932.88 | 482.44 | -964.88 | | | | |
| | Interaction | 8 | -955.4 | -928.74 | 485.7 | -971.4 | 6.523 | 2 | 0.03833 | * |
| **Modularity** | No interaction | 6 | -1194.7 | -1174.7 | 603.33 | -1206.7 | | | | |
| | Interaction | 8 | -1197 | -1170.3 | 606.49 | -1213 | 6.3267 | 2 | 0.04228 | * |
| **F. diversity** | No interaction | 6 | 669.07 | 689.06 | -328.53 | 657.07 | | | | |
| | Interaction | 8 | 670.93 | 697.59 | -327.46 | 654.93 | 2.1392 | 2 | 0.3431 | |
| **F. redundancy** | No interaction | 6 | 957.6 | 977.59 | -472.8 | 945.6 | | | | |
| | Interaction | 8 | 948.25 | 974.92 | -466.13 | 932.25 | 13.344 | 2 | 0.001266 | ** |

**Table S6: Mixed effect model outputs** for all variables tested in this study - fixed effects outputs

| | | Estimate | Std. Error | df | t value | Pr(>\|t\|) | |
|---|---|---|---|---|---|---|---|
| **Bacteria diversity** | (Intercept) | 52.3401 | 2.7689 | 189.6787 | 18.903 | < 2e-16 | *** |
| | DrainageAreaKmLog | 0.9147 | 0.8219 | 189.6787 | 1.113 | 0.267149 | |
| | SeasonSpring | 3.4365 | 3.5615 | 134 | 0.965 | 0.336336 | |
| | SeasonSummer | -5.6958 | 3.5615 | 134 | -1.599 | 0.112123 | |
| | DrainageAreaKmLog:SeasonSpring | 0.2832 | 1.0571 | 134 | 0.268 | 0.789161 | |
| | DrainageAreaKmLog:SeasonSummer | -3.7343 | 1.0571 | 134 | -3.532 | 0.000565 | *** |
| **Fish diversity** | (Intercept) | 0.95731 | 0.20884 | 184.78842 | 4.584 | 8.38E-06 | *** |
| | DrainageAreaKmLog | 0.37095 | 0.06199 | 184.78842 | 5.984 | 1.11E-08 | *** |
| | SeasonSpring | 0.08918 | 0.2626 | 134 | 0.34 | 0.73469 | |
| | SeasonSummer | 0.91455 | 0.2626 | 134 | 3.483 | 0.000671 | *** |
| | DrainageAreaKmLog:SeasonSpring | 0.03738 | 0.07794 | 134 | 0.48 | 0.632311 | |
| | DrainageAreaKmLog:SeasonSummer | -0.31722 | 0.07794 | 134 | -4.07 | 7.99E-05 | *** |
| **Invert diversity** | (Intercept) | 5.97672 | 0.61372 | 200.28743 | 9.739 | < 2e-16 | *** |
| | DrainageAreaKmLog | -0.19869 | 0.18217 | 200.28743 | -1.091 | 0.2767 | |
| | SeasonSpring | 2.54437 | 0.84943 | 134 | 2.995 | 0.00327 | ** |
| | SeasonSummer | 1.12615 | 0.84943 | 134 | 1.326 | 0.18717 | |
| | DrainageAreaKmLog:SeasonSpring | 0.29175 | 0.25213 | 134 | 1.157 | 0.24927 | |
| | DrainageAreaKmLog:SeasonSummer | -0.08574 | 0.25213 | 134 | -0.34 | 0.73434 | |
| **No. Link** | (Intercept) | 333.057 | 35.52 | 186.621 | 9.377 | < 2e-16 | *** |
| | DrainageAreaKmLog | -14.702 | 10.543 | 186.621 | -1.394 | 0.16482 | |
| | SeasonSpring | 130.315 | 45.034 | 134 | 2.894 | 0.00445 | ** |
| | SeasonSummer | 7.172 | 45.034 | 134 | 0.159 | 0.8737 | |
| | DrainageAreaKmLog:SeasonSpring | 27.387 | 13.367 | 134 | 2.049 | 0.04243 | * |
| | DrainageAreaKmLog:SeasonSummer | -12.021 | 13.367 | 134 | -0.899 | 0.37009 | |
| **Link density** | (Intercept) | 5.26487 | 0.41989 | 190.87454 | 12.539 | < 2e-16 | *** |
| | DrainageAreaKmLog | -0.27555 | 0.12463 | 190.87454 | -2.211 | 0.02823 | * |
| | SeasonSpring | 1.51821 | 0.54332 | 133.99999 | 2.794 | 0.00596 | ** |
| | SeasonSummer | 0.22492 | 0.54332 | 133.99999 | 0.414 | 0.67955 | |
| | DrainageAreaKmLog:SeasonSpring | 0.30326 | 0.16127 | 133.99999 | 1.88 | 0.06222 | . |
| | DrainageAreaKmLog:SeasonSummer | 0.05447 | 0.16127 | 133.99999 | 0.338 | 0.73608 | |
| **Nestedness** | (Intercept) | 7.8394 | 0.7652 | 192.7018 | 10.245 | < 2e-16 | *** |
| | DrainageAreaKmLog | -0.6424 | 0.2271 | 192.7018 | -2.829 | 0.00517 | ** |
| | SeasonSpring | 1.5652 | 0.9996 | 134 | 1.566 | 0.11974 | |
| | SeasonSummer | -0.6552 | 0.9996 | 134 | -0.655 | 0.51329 | |
| | DrainageAreaKmLog:SeasonSpring | 0.2129 | 0.2967 | 134 | 0.718 | 0.47419 | |
| | DrainageAreaKmLog:SeasonSummer | 1.0986 | 0.2967 | 134 | 3.703 | 0.00031 | ** |
| **Coherence** | (Intercept) | 0.150699 | 0.013073 | 198.712332 | 11.528 | < 2e-16 | *** |
| | DrainageAreaKmLog | -0.006515 | 0.00388 | 198.712332 | -1.679 | 0.094713 | . |

| | | | | | | | |
|---|---|---|---|---|---|---|---|
| | SeasonSpring | 0.005908 | 0.017773 | 133.999994 | 0.332 | 0.740106 | |
| | SeasonSummer | -0.012769 | 0.017773 | 133.999994 | -0.718 | 0.473709 | |
| | DrainageAreaKmLog:SeasonSpring | 0.006584 | 0.005275 | 133.999994 | 1.248 | 0.214147 | |
| | DrainageAreaKmLog:SeasonSummer | 0.019278 | 0.005275 | 133.999994 | 3.654 | 0.000369 | *** |
| **Connectance** | (Intercept) | 0.089993 | 0.006868 | 199.513706 | 13.103 | < 2e-16 | *** |
| | DrainageAreaKmLog | -0.006097 | 0.002039 | 199.513706 | -2.991 | 0.00313 | ** |
| | SeasonSpring | 0.016533 | 0.009412 | 133.999991 | 1.757 | 0.08127 | . |
| | SeasonSummer | 0.007323 | 0.009412 | 133.999991 | 0.778 | 0.43791 | |
| | DrainageAreaKmLog:SeasonSpring | 0.003207 | 0.002794 | 133.999991 | 1.148 | 0.25299 | |
| | DrainageAreaKmLog:SeasonSummer | 0.008257 | 0.002794 | 133.999991 | 2.956 | 0.00369 | ** |
| **Omnivory** | (Intercept) | 2.14E-02 | 5.93E-03 | 2.01E+02 | 3.604 | 0.000395 | *** |
| | DrainageAreaKmLog | -2.36E-03 | 1.76E-03 | 2.01E+02 | -1.339 | 0.181926 | |
| | SeasonSpring | 9.45E-03 | 8.33E-03 | 1.34E+02 | 1.135 | 0.25838 | |
| | SeasonSummer | 4.60E-03 | 8.33E-03 | 1.34E+02 | 0.553 | 0.581217 | |
| | DrainageAreaKmLog:SeasonSpring | 9.42E-04 | 2.47E-03 | 1.34E+02 | 0.381 | 0.703515 | |
| | DrainageAreaKmLog:SeasonSummer | 5.45E-03 | 2.47E-03 | 1.34E+02 | 2.206 | 0.029077 | * |
| **Func. Diversity** | (Intercept) | 7.12634 | 0.23603 | 192.1954 | 30.193 | <2e-16 | *** |
| | DrainageAreaKmLog | -0.11857 | 0.07006 | 192.1954 | -1.692 | 0.0922 | . |
| | SeasonSpring | 0.55846 | 0.3075 | 134.00323 | 1.816 | 0.0716 | . |
| | SeasonSummer | 0.07407 | 0.3075 | 134.00323 | 0.241 | 0.81 | |
| | DrainageAreaKmLog:SeasonSpring | 0.0899 | 0.09127 | 134.00323 | 0.985 | 0.3264 | |
| | DrainageAreaKmLog:SeasonSummer | -0.03882 | 0.09127 | 134.00323 | -0.425 | 0.6713 | |
| **Func. Redundancy** | (Intercept) | 8.5479 | 0.4583 | 197.4572 | 18.653 | < 2e-16 | *** |
| | DrainageAreaKmLog | 0.3796 | 0.136 | 197.4572 | 2.791 | 0.005773 | ** |
| | SeasonSpring | 0.3478 | 0.6166 | 134 | 0.564 | 0.573655 | |
| | SeasonSummer | -0.7024 | 0.6166 | 134 | -1.139 | 0.256683 | |
| | DrainageAreaKmLog:SeasonSpring | -0.1492 | 0.183 | 134 | -0.815 | 0.416435 | |
| | DrainageAreaKmLog:SeasonSummer | -0.6448 | 0.183 | 134 | -3.523 | 0.000584 | *** |
| **Robustness** | (Intercept) | 0.509513 | 0.004668 | 188.093197 | 109.149 | <2e-16 | *** |
| | DrainageAreaKmLog | -0.004027 | 0.001386 | 188.093197 | -2.906 | 0.0041 | ** |
| | SeasonSpring | -0.010037 | 0.005959 | 134 | -1.684 | 0.0944 | . |
| | SeasonSummer | -0.012496 | 0.005959 | 134 | -2.097 | 0.0379 | * |
| | DrainageAreaKmLog:SeasonSpring | 0.003614 | 0.001769 | 134 | 2.043 | 0.043 | * |
| | DrainageAreaKmLog:SeasonSummer | 0.004135 | 0.001769 | 134 | 2.338 | 0.0209 | * |
| **Modularity** | (Intercept) | 1.03E-02 | 2.57E-03 | 2.00E+02 | 4.026 | 8.06E-05 | *** |
| | DrainageAreaKmLog | -8.44E-05 | 7.62E-04 | 2.00E+02 | -0.111 | 0.9119 | |
| | SeasonSpring | 2.22E-03 | 3.54E-03 | 1.34E+02 | 0.627 | 0.5317 | |
| | SeasonSummer | -1.67E-04 | 3.54E-03 | 1.34E+02 | -0.047 | 0.9624 | |
| | DrainageAreaKmLog:SeasonSpring | 1.91E-04 | 1.05E-03 | 1.34E+02 | 0.182 | 0.856 | |
| | DrainageAreaKmLog:SeasonSummer | 2.37E-03 | 1.05E-03 | 1.34E+02 | 2.257 | 0.0257 | * |

**Table S7: Mixed model outputs** for all variables tested in this study - **Random effect**

| | Groups | Variance | Std.Dev. |
|---|---|---|---|
| **Bacteria diversity** | ID (Intercept) | 34.65 | 5.887 |
| | Residual | 165.93 | 12.881 |
| **Fish diversity** | ID (Intercept) | 0.239 | 0.4889 |
| | Residual | 0.902 | 0.9498 |
| **Invert diversity** | ID (Intercept) | 0.4156 | 0.6447 |
| | Residual | 9.4386 | 3.0722 |
| **No. Link** | ID (Intercept) | 6479 | 80.49 |
| | Residual | 26529 | 162.88 |
| **Link density** | ID (Intercept) | 0.7512 | 0.8667 |
| | Residual | 3.8615 | 1.9651 |
| **Nestedness** | ID (Intercept) | 2.248 | 1.499 |
| | Residual | 13.07 | 3.615 |
| **Coherence** | ID (Intercept) | 0.0003392 | 0.01842 |
| | Residual | 0.004132 | 0.06428 |
| **Connectance** | ID (Intercept) | 7.53E-05 | 0.008679 |
| | Residual | 1.16E-03 | 0.034041 |
| **Omnivory** | ID (Intercept) | 1.41E-05 | 0.003752 |
| | Residual | 9.07E-04 | 0.030109 |
| **F. diversity** | ID (Intercept) | 0.2206 | 0.4696 |
| | Residual | 1.2369 | 1.1122 |
| **F. redundancy** | ID (Intercept) | 0.5204 | 0.7214 |
| | Residual | 4.9738 | 2.2302 |
| **Robustness** | ID (Intercept) | 0.0001056 | 0.01028 |
| | Residual | 0.0004645 | 0.02155 |
| **Modularity** | ID (Intercept) | 8.51E-06 | 0.002917 |
| | Residual | 1.64E-04 | 0.012801 |

**Table S8: Fixed effect analysis of variants for all interaction models.**

| | Sum | Sq | Mean | Sq | NumDF | DenDF | Fvalue | Pr(>F) |
|---|---|---|---|---|---|---|---|---|
| **Bacteria diversity** | DrainageAreaKmLog | 30.42 | 30.42 | 1 | 67 | 0.1834 | 0.6698821 | |
| | Season | 1113.21 | 556.61 | 2 | 134 | 3.3545 | 0.0378858 | * |
| | DrainageAreaKmLog:Season | 2985.99 | 1492.99 | 2 | 134 | 8.9977 | 0.0002155 | *** |
| **Fish diversity** | DrainageAreaKmLog | 38.271 | 38.271 | 1 | 67 | 42.4266 | 1.11E-08 | *** |
| | Season | 13.304 | 6.652 | 2 | 134 | 7.3745 | 0.0009153 | *** |
| | DrainageAreaKmLog:Season | 22.546 | 11.273 | 2 | 134 | 12.497 | 1.06E-05 | *** |
| **Invert diversity** | DrainageAreaKmLog | 13.304 | 13.304 | 1 | 67 | 1.4095 | 0.23933 | |
| | Season | 85.058 | 42.529 | 2 | 134 | 4.5059 | 0.01277 | * |
| | DrainageAreaKmLog:Season | 23.258 | 11.629 | 2 | 134 | 1.2321 | 0.29496 | |
| **No. Link** | DrainageAreaKmLog | 47193 | 47193 | 1 | 67 | 1.7789 | 0.1868 | |
| | Season | 280791 | 140396 | 2 | 134 | 5.2921 | 0.006136 | ** |
| | DrainageAreaKmLog:Season | 242268 | 121134 | 2 | 134 | 4.5661 | 0.012068 | * |
| **Link density** | DrainageAreaKmLog | 13.744 | 13.7442 | 1 | 67 | 3.5593 | 0.06355 | . |
| | Season | 35.129 | 17.5644 | 2 | 134 | 4.5486 | 0.01227 | * |
| | DrainageAreaKmLog:Season | 15.523 | 7.7617 | 2 | 134 | 2.01 | 0.138 | |
| **Nestedness** | DrainageAreaKmLog | 24.758 | 24.758 | 1 | 67 | 1.8943 | 0.1733003 | |
| | Season | 68.101 | 34.05 | 2 | 134 | 2.6052 | 0.0776256 | . |
| | DrainageAreaKmLog:Season | 201.605 | 100.803 | 2 | 134 | 7.7125 | 0.0006755 | *** |
| **Coherence** | DrainageAreaKmLog | 0.003169 | 0.0031692 | 1 | 67 | 0.767 | 0.384278 | |
| | Season | 0.004768 | 0.0023842 | 2 | 134 | 0.577 | 0.562956 | |
| | DrainageAreaKmLog:Season | 0.057028 | 0.0285141 | 2 | 134 | 6.9009 | 0.001404 | ** |
| **Connectance** | DrainageAreaKmLog | 0.0038609 | 0.0038609 | 1 | 67 | 3.3319 | 0.07241 | |
| | Season | 0.003591 | 0.0017955 | 2 | 134 | 1.5495 | 0.21614 | |
| | DrainageAreaKmLog:Season | 0.0102912 | 0.0051456 | 2 | 134 | 4.4405 | 0.01357 | * |
| **Omnivory** | DrainageAreaKmLog | 0.000044 | 0.00004396 | 1 | 67 | 0.0485 | 0.82637 | |
| | Season | 0.0011682 | 0.0005841 | 2 | 134 | 0.6443 | 0.52664 | |
| | DrainageAreaKmLog:Season | 0.0050418 | 0.00252091 | 2 | 134 | 2.7808 | 0.06557 | . |
| **Robustness** | DrainageAreaKmLog | 0.001104 | 0.001104 | 1 | 67 | 2.3767 | 0.12787 | |
| | Season | 0.002293 | 0.0011465 | 2 | 134 | 2.4682 | 0.08858 | . |
| | DrainageAreaKmLog:Season | 0.0030127 | 0.0015064 | 2 | 134 | 3.243 | 0.04213 | * |
| **Modularity** | DrainageAreaKmLog | 0.00045633 | 0.00045633 | 1 | 67 | 2.7849 | 0.09982 | . |
| | Season | 0.00009284 | 0.00004642 | 2 | 134 | 0.2833 | 0.75374 | |
| | DrainageAreaKmLog:Season | 0.00103004 | 0.00051502 | 2 | 134 | 3.1432 | 0.04634 | * |
| **F. Diversity** | DrainageAreaKmLog | 5.9845 | 5.9845 | 1 | 66.999 | 4.8383 | 0.03129 | * |
| | Season | 4.8138 | 2.4069 | 2 | 134.003 | 1.9459 | 0.14687 | |
| | DrainageAreaKmLog:Season | 2.5894 | 1.2947 | 2 | 134.003 | 1.0467 | 0.35395 | |
| **F. Redundancy** | DrainageAreaKmLog | 8.961 | 8.961 | 1 | 67 | 1.8016 | 0.184056 | |
| | Season | 14.977 | 7.488 | 2 | 134 | 1.5055 | 0.225627 | |
| | DrainageAreaKmLog:Season | 67.664 | 33.832 | 2 | 134 | 6.8021 | 0.001536 | ** |

**Table S9: Contrast testing output from emtrends().** The pairwise comparison of the mixed effect model slopes

| | contrast | estimate | SE | df | t.ratio | p.value | |
|---|---|---|---|---|---|---|---|
| **Bacteria diversity** | Spring - Summer | 4.018 | 1.06 | 134 | 3.8 | 0.0006 | *** |
| | Spring - Autumn | 0.283 | 1.06 | 134 | 0.268 | 0.9612 | |
| | Summer - Autumn | -3.734 | 1.06 | 134 | -3.532 | 0.0016 | ** |
| **Fish diversity** | Spring - Summer | 0.3546 | 0.0779 | 134 | 4.549 | <.0001 | **** |
| | Spring - Autumn | 0.0374 | 0.0779 | 134 | 0.48 | 0.8811 | |
| | Summer - Autumn | -0.3172 | 0.0779 | 134 | -4.07 | 0.0002 | *** |
| **Invert diversity** | Spring - Summer | 0.3775 | 0.252 | 134 | 1.497 | 0.2955 | |
| | Spring - Autumn | 0.2918 | 0.252 | 134 | 1.157 | 0.4809 | |
| | Summer - Autumn | -0.0857 | 0.252 | 134 | -0.34 | 0.9383 | |
| **No. Link** | Spring - Summer | 39.4 | 13.4 | 134 | 2.948 | 0.0105 | |
| | Spring - Autumn | 27.4 | 13.4 | 134 | 2.049 | 0.1047 | |
| | Summer - Autumn | -12 | 13.4 | 134 | -0.899 | 0.6417 | |
| **Link density** | Spring - Summer | 0.2488 | 0.161 | 134 | 1.543 | 0.2744 | |
| | Spring - Autumn | 0.3033 | 0.161 | 134 | 1.88 | 0.1483 | |
| | Summer - Autumn | 0.0545 | 0.161 | 134 | 0.338 | 0.9391 | |
| **Nestedness** | Spring - Summer | -0.886 | 0.297 | 134 | -2.985 | 0.0094 | ** |
| | Spring - Autumn | 0.213 | 0.297 | 134 | 0.718 | 0.7535 | |
| | Summer - Autumn | 1.099 | 0.297 | 134 | 3.703 | 0.0009 | *** |
| **Coherence** | Spring - Summer | -0.01269 | 0.00528 | 134 | -2.406 | 0.0457 | ** |
| | Spring - Autumn | 0.00658 | 0.00528 | 134 | 1.248 | 0.4271 | |
| | Summer - Autumn | 0.01928 | 0.00528 | 134 | 3.654 | 0.0011 | ** |
| **Connectance** | Spring - Summer | -0.00505 | 0.00279 | 134 | -1.808 | 0.171 | |
| | Spring - Autumn | 0.00321 | 0.00279 | 134 | 1.148 | 0.4864 | |
| | Summer - Autumn | 0.00826 | 0.00279 | 134 | 2.956 | 0.0102 | ** |
| **Omnivory** | Spring - Summer | -0.004509 | 0.00247 | 134 | -1.825 | 0.1654 | |
| | Spring - Autumn | 0.000942 | 0.00247 | 134 | 0.381 | 0.923 | |
| | Summer - Autumn | 0.005451 | 0.00247 | 134 | 2.206 | 0.0738 | |
| **Func. Diversity** | Spring - Summer | 0.1287 | 0.0913 | 134 | 1.41 | 0.3385 | |
| | Spring - Autumn | 0.0899 | 0.0913 | 134 | 0.985 | 0.5876 | |
| | Summer - Autumn | -0.0388 | 0.0913 | 134 | -0.425 | 0.9052 | |
| **Func. Redundancy** | Spring - Summer | 0.496 | 0.183 | 134 | 2.708 | 0.0208 | ** |
| | Spring - Autumn | -0.149 | 0.183 | 134 | -0.815 | 0.6943 | |
| | Summer - Autumn | -0.645 | 0.183 | 134 | -3.523 | 0.0017 | ** |
| **Robustness** | Spring - Summer | -0.000521 | 0.00177 | 134 | -0.295 | 0.9533 | |
| | Spring - Autumn | 0.003614 | 0.00177 | 134 | 2.043 | 0.1059 | |
| | Summer - Autumn | 0.004135 | 0.00177 | 134 | 2.338 | 0.054 | |
| **Modularity** | Spring - Summer | -0.002179 | 0.00105 | 134 | -2.075 | 0.099 | |
| | Spring - Autumn | 0.000191 | 0.00105 | 134 | 0.182 | 0.9819 | |
| | Summer - Autumn | 0.002371 | 0.00105 | 134 | 2.257 | 0.0656 | |

**Table S10: Contrast testing output from emmeans**. The pairwise comparison of means between seasons for all variables included in the analysis

| | contrast | estimate | SE | df | t.ratio | p.value | |
|---|---|---|---|---|---|---|---|
| **Bacteria diversity** | Spring - Summer | 19.8 | 2.19 | 134 | 9.027 | <.0001 | **** |
| | Spring - Autumn | 4.19 | 2.19 | 134 | 1.91 | 0.1398 | |
| | Summer - Autumn | -15.61 | 2.19 | 134 | -7.117 | <.0001 | **** |
| **Fish diversity** | Spring - Summer | 0.1159 | 0.162 | 134 | 0.717 | 0.7539 | |
| | Spring - Autumn | 0.1884 | 0.162 | 134 | 1.165 | 0.476 | |
| | Summer - Autumn | 0.0725 | 0.162 | 134 | 0.448 | 0.8953 | |
| **Invert diversity** | Spring - Summer | 2.42 | 0.523 | 134 | 4.627 | <.0001 | **** |
| | Spring - Autumn | 3.319 | 0.523 | 134 | 6.345 | <.0001 | **** |
| | Summer - Autumn | 0.899 | 0.523 | 134 | 1.718 | 0.2023 | |
| **No. Link** | Spring - Summer | 227.8 | 27.7 | 134 | 8.213 | <.0001 | **** |
| | Spring - Autumn | 203 | 27.7 | 134 | 7.321 | <.0001 | **** |
| | Summer - Autumn | -24.7 | 27.7 | 134 | -0.892 | 0.6462 | |
| **Link density** | Spring - Summer | 1.95 | 0.335 | 134 | 5.84 | <.0001 | **** |
| | Spring - Autumn | 2.32 | 0.335 | 134 | 6.944 | <.0001 | **** |
| | Summer - Autumn | 0.37 | 0.335 | 134 | 1.104 | 0.513 | |
| **Nestedness** | Spring - Summer | -0.131 | 0.615 | 134 | -0.212 | 0.9754 | |
| | Spring - Autumn | 2.13 | 0.615 | 134 | 3.461 | 0.0021 | ** |
| | Summer - Autumn | 2.261 | 0.615 | 134 | 3.674 | 0.001 | *** |
| **Coherence** | Spring - Summer | -0.015 | 0.0109 | 134 | -1.372 | 0.3583 | |
| | Spring - Autumn | 0.0234 | 0.0109 | 134 | 2.137 | 0.0863 | |
| | Summer - Autumn | 0.0384 | 0.0109 | 134 | 3.509 | 0.0018 | *** |
| **Connectance** | Spring - Summer | -0.0042 | 0.0058 | 134 | -0.724 | 0.7498 | |
| | Spring - Autumn | 0.025 | 0.0058 | 134 | 4.322 | 0.0001 | *** |
| | Summer - Autumn | 0.0292 | 0.0058 | 134 | 5.046 | <.0001 | **** |
| **Omnivory** | Spring - Summer | -0.00712 | 0.00513 | 134 | -1.39 | 0.3492 | |
| | Spring - Autumn | 0.01195 | 0.00513 | 134 | 2.331 | 0.0549 | |
| | Summer - Autumn | 0.01907 | 0.00513 | 134 | 3.721 | 0.0008 | *** |
| **Func. Diversity** | Spring - Summer | 0.826 | 0.189 | 134 | 4.363 | 0.0001 | *** |
| | Spring - Autumn | 0.797 | 0.189 | 134 | 4.21 | 0.0001 | *** |
| | Summer - Autumn | -0.029 | 0.189 | 134 | -0.153 | 0.9872 | |
| **Func. Redundancy** | Spring - Summer | 2.3658 | 0.38 | 134 | 6.231 | <.0001 | **** |
| | Spring - Autumn | -0.0482 | 0.38 | 134 | -0.127 | 0.9911 | |
| | Summer - Autumn | -2.414 | 0.38 | 134 | -6.358 | <.0001 | **** |
| **Robustness** | Spring - Summer | 0.001075 | 0.00367 | 134 | 0.293 | 0.9538 | |
| | Spring - Autumn | -0.000443 | 0.00367 | 134 | -0.121 | 0.992 | |
| | Summer - Autumn | -0.001518 | 0.00367 | 134 | -0.414 | 0.9101 | |
| **Modularity** | Spring - Summer | -0.0034 | 0.00218 | 134 | -1.56 | 0.2667 | |
| | Spring - Autumn | 0.00273 | 0.00218 | 134 | 1.251 | 0.4255 | |
| | Summer - Autumn | 0.00613 | 0.00218 | 134 | 2.811 | 0.0156 | * |

**Table S11: β-diversity against river distance for each group and each season**. β-diversity is calculated based on Jaccard dissimilarity where 0 would indicate identical communities. β diversity was further separated into genus loss (Nestedness) and replacement (Turnover). Correlations reported are based on Mantel statistic and p-values (significant p-values are highlighted in bold)

| Group | Season | Jaccard dissimilarity | Taxon loss (Nestedness) | Taxon replacement (Turnover) |
|---|---|---|---|---|
| Fish | Spring | 0.1434<br>**p = 0.002** | 0.2187<br>**p = 0.001** | 0.07043<br>p = 0.891 |
| | Summer | 0.1712<br>**p = 0.001** | -0.08374<br>p = 0.989 | 0.2031<br>**p = 0.001** |
| | Autumn | 0.3214<br>**p = 0.001** | 0.0587<br>**p = 0.05** | 0.2691<br>**p = 0.001** |
| Invertebrate | Spring | 0.0952<br>**p = 0.031** | -0.00215<br>p = 0.464 | 0.06638<br>p = 0.051 |
| | Summer | 0.1685<br>**p = 0.001** | -0.07856<br>p = 0.976 | 0.142<br>**p = 0.002** |
| | Autumn | 0.1114<br>**p = 0.013** | -0.01447<br>p = 0.597 | 0.07066<br>**p = 0.047** |
| Bacteria | Spring | -0.05822<br>p = 0.834 | -0.0655<br>p = 0.928 | 0.01599<br>p = 0.34 |
| | Summer | **0.1785**<br>**p = 0.003** | 0.08859<br>**p = 0.044** | 0.06969<br>p = 0.068 |
| | Autumn | 0.06779<br>p = 0.12 | -0.04308<br>p = 0.747 | 0.1185<br>**p = 0.014** |

**Table S12: Primer selection for library preparation:** 12S and COI primers included a modification to include the Nextera® transposase sequences in the first PCR, whereas 16S included this modification in the second PCR.

| Marker | Fragment | Forward primer | Reverse primer | Reference |
|---|---|---|---|---|
| **12S** | **106 bp** | 5’-TACTGGGATTAGATACCCC-3’ | 5’-CTAGAACAGGCTCCTCTAG-3 | [14] |
| **COI** | **313 bp** | mICOIintF: 5’-GGWACWGGWTGAACWGTWTAYCCYCC-3’ | jgHCO2198: 5’-TAIACYTCIGGRTGICCRAARAAYCA-3’ | [15, 16] |
| **16S** | **450 bp** | U341F: 5’-CCTACGGGDGGCWGCA-3’ | U806R: 5’GACTACHVGGGTMTCTAATC-3’ | [17] |

**Table S13: Positive sample information.** For each library a positive control was included to be used in determining possible contamination within the library preparation.

| Marker | Type | Sequence if applicable |
|---|---|---|
| **12S** | **Tissue extraction (*Gadus morhua*)** | TACNTNNTGTTATGGTTCCGTTTAACATTGATG<br>GTTTTATTACCCAAACCATNCCGCCTGGGAAC<br>TACGAGCAATAGCTTAAAACCCAAAGGACTTN<br>GGCGGTGCTTTAGACCCCCCTAGAGGAGCCTG<br>TTCTAGCTGTCTCTTATACACATCTCCGGCCCA<br>CGAGACNGAGCTAGAACNAGCTCCTCTNGNGG<br>GTNTAANGCCCGCCANTCCTTTGGGTTTTAAG<br>CTATTGCTCGTANNTCCAGCGATGGTTTGGGTA<br>TAAAACATCAATGTTAACNACCATACTNNTGG<br>GGTATCTAATCCAGTACTGTCTCTTATACCGTC<br>TACGCTCCNCANCCCTGTTAGACGCATATTGAA<br>TGNGATCGCCGGTCGGGGTATGAAAAACAACG<br>GCANCNCCGGNCAANACATCACACCCGCCNGG<br>AAAAACCAAACCNGCNTTTNTAAACCTGCTTGC<br>ACAGGCCGTNCGCGTCATTCTTCACGGNGNCGC<br>ATGTTTTNGTTTCGNCATTTTGCCGCCTGCGTGG<br>GCTGCGTGAGCAGCACACCCAGCCAGCCCCTTN<br>NCTGCCGTTGNCGGNGNCAANNGCNTCNGGACA<br>GCACTCCNGCNTGNCTCGTCTAGCACCACCACA<br>CNACAGCANAGAGCNGATGCTCNCCGCCAACGA<br>AGCCAGGNNGNTGNTTGAGATNCNTAAGTATNA<br>CCTACTGGCGCNNNATCTTGAGATNCGCGTCCA<br>AGTCTAGCCNATAATNNGCAAGATAGNTTGTGT<br>CNAGGCNGTCATCGACNTCAGCAACGCTGNGNT<br>TANNGNTNGTTG |
| **COI** | **Synthetic sequence** | GGAACAGGTTGAACTGTATATCCCCCATCAACC<br>TAGTTACGAAGAGCTATAGATCATATAATCCTT<br>AAGTGGAATGTTAATGTGAGTTCAATATGATAC<br>ACGCCACAGACTCATGTATGTGGATCGGAAGCC<br>AGCTGTTTCCGACCTCGGAGCCGAGAGTGGTTT<br>CTGAATTACACATGTAAGATAAAATCATTAAAG<br>GTACTAACTCACGAAACCTCAGGATATGCGTGG<br>TTTGCTGAGATTTCTATTTTCTCGTTCTTGATTTA<br>ACCACGTAAAATGTGTGAAAACTAAAGGTTCTA<br>GCATTTCTAAGGATCACTACGCCTAACGTCTCAC<br>TTTATCTTAAATTTGATTTTTTGGTCACCCTGAAGT<br>TTA |
| **16S** | **Bacteria stock community** | *Bacillus subtilis, Bacillus brevis* and *Serratia fonticuli* |

### Methods:

#### First PCR for library preparation

*12S - Vertebrate library*

The first PCR was carried out in a total volume of 25 µL containing: 0.5 µM each primer, 0.4 mg/mL BSA (New England Biolabs, Ipswich, MA, USA), 12.5 µL Q5® HighFidelity 2X Master Mix (New England Biolabs), and 2 µL of DNA template per reaction. PCR profiles were as follows: initial denaturation 98 °C for 5 min followed by 35 cycles of 98 °C for 10 s, 58 °C for 20 s, and 72 °C for 30 s, and a final extension step of 72 °C for 7 min, before the plates were cooled down to 10 °C.

*COI - Metazoan library*

The first PCR was carried out in a total volume of 25 µL containing: 0.5 µM each of each primer, AmpliTaq Gold 360° (1.25 U/µL), 1x Buffer I (Thermo Fisher Scientific, MD, USA), 0.1 mg/µL BSA, 0.2 mM dNTP, 1 mM $MgCl_2$, SigmaFree water and 2 µL of DNA template per reaction. PCR profiles were carried out using a touchdown protocol as follows: initial denaturation at 95 °C for 10 min, the first 25 cycles started with the denaturation at 95 °C for 15 s, annealing at 62 °C for 30 s, followed by extension at 72 °C for 30 s. After this the cycler performed 16 cycles where the annealing temperature was reduced by one degree each cycle, performing the last cycle at a temperature of 45 degrees. Final extension was performed at 72 °C for 5 min before the plates were cooled down to 10 °C.

*16S – Archaea and Bacteria library*

Library preparation for the 16S V3-V4 region followed a three-step PCR library preparation using the method described in [18] with minor modifications. In brief, the first PCR was performed in 15 µL volumes containing: 0.5 µM of each forward and reverse primer, 1x supplied buffer (Faststart TAQ, Roche, Inc., Basel, Switzerland), 1 mg/µL BSA, 0.18 mM dNTPs, 2.0 mM $MgCl_2$, 0.05 units per µL Taq DNA polymerase (Faststart TAQ, Roche, Inc.) and 2 µL of DNA template per reaction. This was carried out in triplicate and then pooled. Each plate was then cleaned using Exo I Nuclease (EXO I) and Shrimp Alkaline Phosphatase (SAP) (Thermo Fisher Scientific Inc., Waltham, Maryland USA). The master mix consisted of 1.6 U/µL Exo I and 0.15 U/µL SAP in a total volume of 1.1 µL which was then added to 7.5 µL of the PCR product. Products were heated to 37 °C for 15 minutes, followed by 15minutes at 80 °C, and was then cooled to 4 °C and stored at –20 °C. The second PCR was conducted with the same PCR conditions as the first PCR except the forward and reverse primers were modified to include the Nextera® transposase sequences (Microsynth, AG, Balgach, Switzerland) and only 1 µL of cleaned PCR product was used in the reaction.

**Data preparation workflow - steps and parameters:**

*12S*

**Step A - Data Quality Check:** usearch v11.0.667

**Step B - Trimming and Merging:** usearch v11.0.667

Trim: R1:30nt, R2:40nt

Merge: Min. overlap: 15 bp, Max difference: 10, Min Identity (%): 70, Min. Merged length: 50 and Min. merged quality: 0.

**Step C - Trim Full-Length Primer Sites:** usearch v11.0.667

Amplicon range: 100-300

No. of mismatches: 1

Coverage: full-length

**Step D: Size selection and quality filtering:** PRINSEQ-lite 0.20.4

Size Range: 100-250

GC Range: 30-70

Min Q Mean: 20

**Step E - Clustering - UNOISE for ZOTUs:** usearch v11.0.667

Min Abundance Size: 10

**Step F - Taxonomic Assignment:** SINTAXv11.0.667

Confidence threshold: 0.85

Reference: NCBI BLAST[19] based reference (v200416)

*COI (Metazoan)*

**Step A - Data Quality Check:** usearch v11.0.667

**Step B - Trimming and Merging:** usearch v11.0.667

Trim: R1: 40nt, R2: 50nt

Merge: Min. overlap: 40 bp, Min Identity (%): 70, Min. Merged length: 100

**Step C - Trim Full-Length Primer Sites:** usearch v11.0.667

Amplicon range: 100-600

No. of mismatches: 1

Coverage: full-length

**Step D: Size selection and quality filtering:** PRINSEQ-lite 0.20.4

Size Range: 300-450

GC Range: 30-70

Min Q Mean: 20

**Step E - Clustering - UNOISE for ZOTUs:** usearch v11.0.667

**Step F - Taxonomic Assignment Predictions:** SINTAX v11.0.667

Confidence threshold: 0.85

Reference: Custom made reference database consisting of MIDORI[20] (v20180221) and EPT (v200420). Supplemented with unpublished macroinvertebrate sequences.

*16S (Archaea and Bacteria)*

**Step A - Data Quality Check:** usearch v11.0.667

**Step B - Trimming and Merging:** usearch v11.0.667 and FLASH v1.2.11

Trim: R1: 20nt, R2: 50nt

Merge: Min. overlap: 15 bp, Max. overlap: 300

**Step C - Trim Full-Length Primer Sites:** usearch v11.0.667

Amplicon range: 100-2000

No. of mismatches: 1

Coverage: full-length

**Step D: Size selection and quality filtering:** PRINSEQ-lite 0.20.4

Size Range: 300-500

GC Range: 30-70

Min Q Mean: 20

Low complexity filter: dust (30)

**Step E - Clustering – UNOISE3 for ZOTUs:** usearch v11.0.667

**Step F - Taxonomic Assignment Predictions:** SINTAX v11.0.667

Confidence threshold: 0.85

References: SILVA[21] (V128)
